# Structural and functional sex differences in the ventral pallidal vasopressin system are associated with the sex-specific regulation of juvenile social play behavior in rats

**DOI:** 10.1101/2021.01.31.429043

**Authors:** J.D.A. Lee, C.J. Reppucci, S.M. Bowden, E.D.M. Huez, R. Bredewold, A.H. Veenema

**Author notes:** **Corresponding Author:** Jessica D.A. Lee, M.A., Department of Psychology. Michigan State University, Interdisciplinary Science and Technology Building 4401, 766 Service Rd, East Lansing, MI, 48824.

## Abstract

The ventral pallidum (VP) has been implicated in the regulation of rewarding adult social behaviors, such as pair-bonding and sociosexual motivation. However, the role of the VP in regulating rewarding juvenile social behaviors, such as social play, is unknown. Social play is predominantly displayed by juveniles of many mammalian species and engagement in social play helps develop social competence. In this study, we determined whether the VP is involved in regulating social play in juvenile rats by temporarily inactivating the VP via bilateral infusions of muscimol, the GABA_A_ receptor agonist. Muscimol treatment decreased social play duration in males and females compared to the same-sex control groups. We then focused on the vasopressin (AVP) system in the VP as one potential modulator of social play. We examined the organization of the AVP system in the VP in juvenile rats and found robust sex differences, with denser AVP-immunoreacive fibers and denser vasopressin 1a receptor (V1aR) binding in males compared to females, but a greater number of V1aR-expressing cells in females compared to males. Next, we determined whether exposure to social play changed the activation of V1aR-expressing VP cells in male and female juvenile rats. We found that exposure to social play enhanced the number of V1aR-expressing VP cells co-expressing *fos*, a marker of neuronal activation, in males only. Finally, we determined the causal involvement of AVP signaling in the VP in social play behavior by infusion of a specific V1aR antagonist into the VP prior to social play exposure. We found that V1aR blockade in the VP increased social play duration in juvenile male rats but decreased social play duration in juvenile female rats compared to same-sex control groups. These findings reveal structural and functional sex differences in the AVP system in the VP that are associated with the sex-specific regulation of juvenile social play behavior.

## Introduction

The Social Decision-Making Network (SDMN) consists of two well-known brain systems, specifically the mesocorticolimbic reward system (Kelley and Berridge, 2002) and the Social Behavior Network (Newman, 1999), that together function to ensure the coordination of appropriate social behavioral responses (O’Connell and Hofmann, 2011). One brain structure in the SDMN that is a component of the mesocorticolimbic reward system, and which regulates diverse rewarding and motivating behaviors, is the ventral pallidum (VP; Smith and Berridge, 2005; O’Connell and Hofmann, 2011; Berridge and Kringelbach, 2015). The VP has been implicated in the regulation of rewarding social behaviors, including partner preference in adult prairie voles (Pitkow et al., 2001; Lim and Young, 2004; Lim et al., 2004b) and sociosexual motivation in adult rats (DiBenedictis et al., 2020). Notably, both lines of work indicate an important role of arginine vasopressin (AVP) signaling in the VP for the regulation of these social behaviors. For example, blockade of AVP signaling via local bilateral infusion of the vasopressin 1a receptor (V1aR) antagonist in the VP prevented partner preference in monogamous adult male prairie voles (Pitkow et al., 2001; Lim and Young, 2004), while viral vector-mediated overexpression of V1aR in the VP induced a partner preference in non-monogamous adult male meadow voles (Lim et al., 2004b). Furthermore, blockade of AVP signaling in the VP decreased investigation of the opposite sex in adult male rats, while it increased investigation of the opposite sex in adult female rats (DiBenedictis et al., 2020). This latter study highlights a sex-specific role of AVP signaling in the VP in the regulation of adult social behaviors. However, it is unclear whether intra-VP AVP signaling is also important for the regulation of rewarding social behaviors at younger ages, such as social play behavior in juveniles, and if so, whether it does this in sex-specific ways.

Social play behavior (also known as play-fighting and rough-and-tumble play; Pellis and Pellis, 1987; Pellis and Iwaniuk, 2000) is predominantly displayed by juveniles of various mammalian species. Engagement in social play behavior is essential for the development of social competence later in adulthood (Bekoff, 1974; Taylor, 1980; van den Berg et al., 1999; Nijhof et al., 2018). Recent studies demonstrated the involvement of AVP signaling within the brain in the regulation of social play behavior of male and female juvenile rats (Veenema et al., 2013; Bredewold et al., 2014; Paul et al., 2014; Lukas and Wöhr, 2015). For example, administration of an AVP V1a receptor antagonist into the lateral septum (LS) increased social play duration in juvenile males but decreased social play duration in juvenile females (Veenema et al., 2013; Bredewold et al., 2014). In addition, AVP mRNA expression in the bed nucleus of the stria terminalis (BNST) negatively correlated with social play behavior in juvenile males, but not juvenile females (Paul et al., 2014). Because AVP neurons in the BNST project to the LS (De Vries and Buijs, 1983), these findings together suggest that the BNST is a likely source of AVP’s action in the LS where AVP exerts an inhibitory effect on social play in males and an excitatory effect on social play in females. It was recently shown in adult rats that AVP neurons in the BNST also project to the VP (DiBenedictis et al., 2020). However, whether AVP signaling in the VP regulates social play in a similar sex-specific way as it does in the LS is unclear.

In the present study, we aimed to determine the role of the VP, as well as of intra-VP AVP signaling, in the regulation of social play behavior in male and female juvenile rats. To this end, we first determined whether overall VP activity was necessary for the expression of social play behavior through temporary inactivation of the VP by infusing the GABA_A_ receptor agonist muscimol into the VP. Second, we examined the organization of the AVP system in the VP through analysis of AVP-immunoreactive (AVP-ir) fibers, V1aR binding density, and the number of V1aR-expressing cells. Third, we determined whether exposure to social play altered the activation of the VP and of V1aR-expressing cells in the VP using *fos* as an indirect marker of neuronal activity (Morgan and Curran, 1991). Lastly, we determined the role of intra-VP AVP signaling in the expression of social play behavior through infusion of a specific V1aR antagonist or synthetic AVP into the VP. We hypothesized that VP activation is critical for the expression of social play behavior in juvenile rats and that exposure to social play enhances recruitment of VP cells. Furthermore, we hypothesized that intra-AVP signaling in the VP regulates social play behavior in a sex-specific manner and that exposure to social play recruits VP V1aR-expressing cells differently in male and female juvenile rats.

## Methods

### Subjects

Three-week-old male and female Wistar rats were obtained from Charles River Laboratories (Raleigh, NC) and maintained under standard laboratory conditions (12 h light/dark cycle, lights off at 14:00 h, food and water available *ad libitum*). Rats were housed in single-sex groups of four in standard rat cages (48 × 27 × 20 cm or 41.2 × 30 × 23.4 × cm) unless otherwise mentioned. The experiments were conducted in accordance with the National Institute of Health *Guidelines for Care and Use of Laboratory Animals* and approved by the Boston College and Michigan State University Institutional Animal Care and Use Committees.

### Experiment 1: Determine the effect of ventral pallidal infusions of the GABA_A_ receptor agonist muscimol on social play behavior

In order to determine whether VP activity is necessary for the expression of social play behavior, temporary inactivation of the VP was induced via bilateral infusions of the GABA_A_ receptor agonist muscimol into the VP. Juvenile (31 days old) male and female rats underwent stereotaxic surgery after one week of handling. During surgery, experimental subjects were maintained under isoflurane anesthesia, 2-4% as needed (Henry Schein, Melville, NY). Guide cannulae (22 gauge, 9mm; Plastics One, Roanoke, VA) were bilaterally implanted 2 mm dorsal to the VP. Coordinates for the VP from bregma were: 0.25 mm rostral to bregma, 2.4 mm lateral to the midline, and 6.0 mm ventral to the surface of the skull (Paxinos and Watson, 2007). Cannulae were implanted under an angle of 10° from the midsagittal plane to avoid damage to the sagittal sinus. Cannulae were fixed to the skull with four stainless steel screws and dental cement and closed with a dummy cannula (Plastics One, Roanoke, VA). Experimental subjects were given an s.c. injection of Rimadyl (Covetrus 10000319; 10 mg/kg) immediately after surgery and once a day for an additional two days after surgery. Immediately after surgery, experimental subjects were individually housed in standard rat cages until the end of the experiment. Experimental subjects’ body weights were monitored for three days post surgery to ensure normal recovery.

Four days after surgery, experimental subjects (35 days old) received bilateral infusions of either vehicle (aCSF; pH 7.4; males n = 5; females n = 4) or the GABA_A_ receptor agonist muscimol (Sigma Aldrich, M1523; 10ng/side, males n = 8; females n = 7) into the VP 20 min prior to exposure to the social play test (described below). The infusions (0.5uL/side) were given over the course of 45 s via an injector cannula (28 gauge; Plastics One, Roanoke, VA) that extended 2 mm beyond the guide cannula and was connected via polyethylene tubing to a 2 μL syringe (Hamilton Company #88400) mounted onto a microinfusion pump (GenieTouch, Kent Scientific, Torrington, CT). The injector cannula was kept in place for an additional 30 s following infusion to allow for tissue uptake before being replaced by the dummy cannula. Time of administration (Veenema et al., 2013) and drug concentration (Numan et al., 2005) were based on previous studies showing changes in social behaviors of rats upon intracerebral drug infusions.

Social play was assessed in 35 day-old juvenile rats because social play is at its peak at this age (Panksepp, 1981; Pellis and Pellis, 1987; Paul et al., 2014). Social play testing was performed according to Veenema and Neumann (2009). Briefly, testing was performed during the first hour of the dark phase in which the experimental subject’s homecage was removed from the cage rack, the wire cage lid was removed and replaced with a Plexiglass lid. A tripod and video camera were set up above each cage to record the tests. An unfamiliar sex- and age-matched stimulus subject was then placed in the cage. The social play test lasted 10 minutes, during which time the experimental subject was allowed to freely interact with the stimulus subject. Stimulus subjects were striped with a permanent marker 30–60 min prior to social play testing in order to distinguish between the experimental and stimulus subjects during later video analysis. Food and water were not available during the 10 min tests but were immediately returned upon completion of each session.

Social play tests were videotaped and behavior was measured by a researcher blind to the sex and drug treatment conditions of the experimental subjects using SolomonCoder software (https://solomon.andraspeter.com/). The following behaviors were scored for the experimental subjects according to Veenema and Neumann (2009): duration of social play (the total amount of time spent in playful social interactions including nape attacks, pinning, and supine poses), duration of social investigation (the experimental subject is sniffing the anogenital and head/neck regions of the stimulus subject), duration of allogrooming (the experimental subject is grooming the stimulus subject), duration of non-social cage exploration (the experimental subject is walking, rearing, sitting, or engaging in other neutral behaviors), number of nape attacks (the experimental subject displays nose attacks or nose contacts toward the nape of the neck of the stimulus subject), number of pins (the experimental subject holds the stimulus subject on its back in a supine position), and number of supine poses (the experimental subject is pinned by the stimulus subject).

At the end of the experiment, experimental subjects were euthanized with CO_2_ and charcoal was injected as a marker to check proper placement of the cannulae. Brains were extracted, rapidly frozen in methylbutane cooled in dry ice, and stored at −80°C. Brains were sliced in 30 μm sections on the cryostat (Leica CM3050 S) and every third section was mounted directly on Superfrost Plus Slides (Fisher Scientific). Sections were stained with thionin and coverslipped with Permount mounting medium (SP15-100, Fisher Scientific, Waltham, MA). Slides were examined using light microscopy and cannula placements were mapped using The Rat Brain Atlas of Paxinos and Watson (2007). Only rats with bilateral cannulae tracks terminating in the VP were included in statistical analyses (Fig. 1).

**Figure 1.**
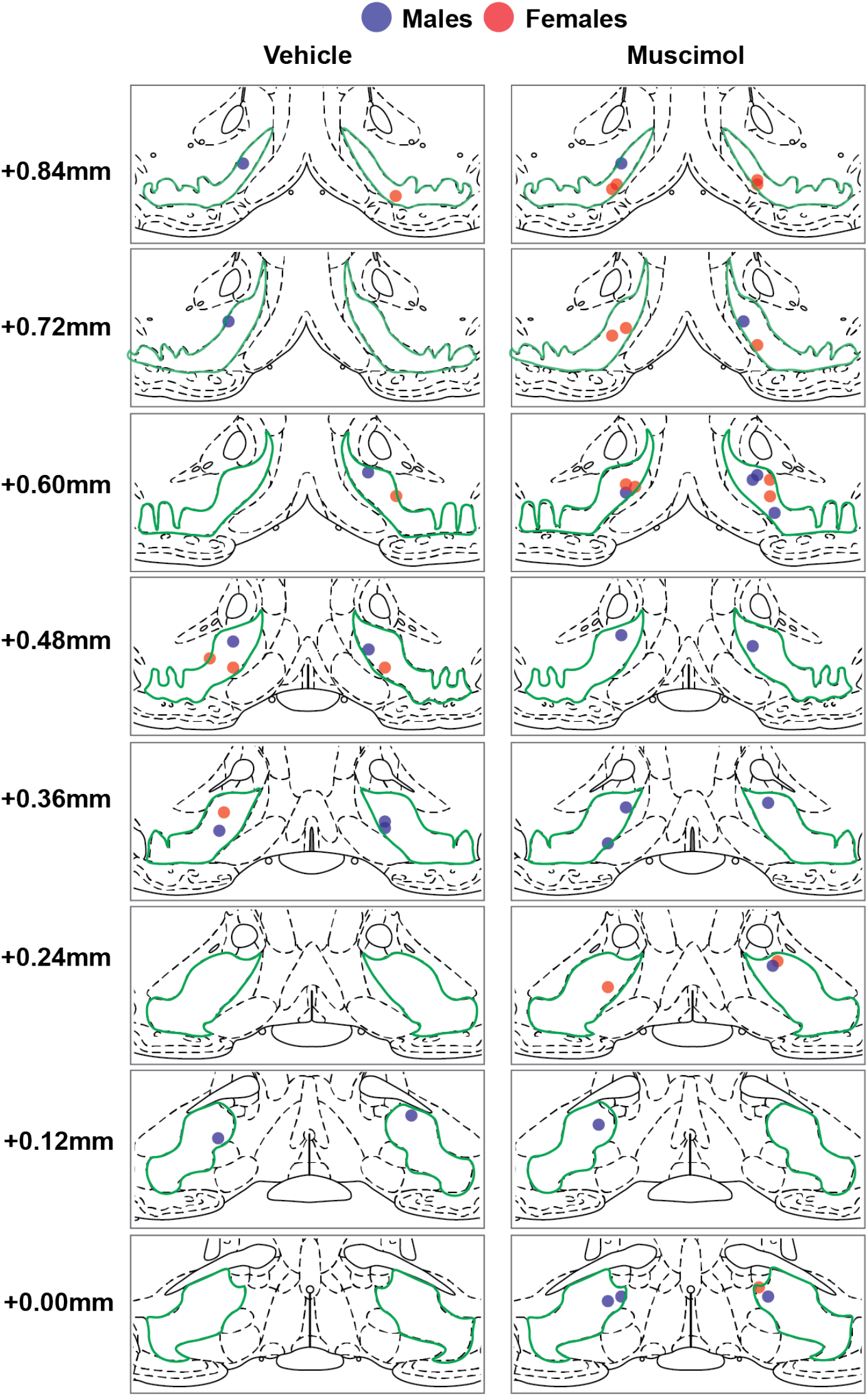
Bilateral cannulae placements in the ventral pallidum (based on the central placement of charcoal that was injected as marker) for Experiment 1 shown on modified rat brain atlas templates (Paxinos and Watson, 2007). Ventral pallidum is outlined in green; numbers to the left of the atlas templates represent distance from Bregma.

### Experiment 2: Determine vasopressin fiber density across the anterior to posterior axis of the ventral pallidum

To test for potential sex differences in AVP fiber density in the VP, AVP-immunoreactive (AVP-ir) fiber density was measured across the anterior-posterior axis of the VP. Juvenile (36 days old) male (n = 4) and female (n = 4) rats were deeply anesthetized with isoflurane before being transcardially perfused with 0.9 % saline followed by 4% paraformaldehyde in 0.1 M borate buffer (pH 9.5). Brains were then extracted and post-fixed for 24 hours in 4% paraformaldehyde in 0.1 M borate buffer. Brains were then cryoprotected in 30% sucrose for 48 hours. Afterwards, brains were rapidly frozen in methylbutane, and stored at −45°C until further histological processing. Brains were sliced in 30 μm coronal sections into 3 series on a cryostat (Leica CM3050 S) and stored free-floating in antifreeze cryoprotectant [0.25 mM polyvinylpyrolidone, 30% sucrose, 30% ethylene glycol in tris-buffered saline (50mM, pH 7.6; TBS)] at −20°C.

AVP-immunoreactivity was visualized using a monoclonal primary antibody provided by Dr. Harold Gainer (NINDS). These highly specific antibodies were raised against mammalian AVP-associated neurophysins and exhibit no cross reactivity (Ben-Barak et al., 1985). AVP immunohistochemistry was performed according to DiBenedictis et al. (2017). Briefly, one series of tissue sections was first washed in TBS (50mM, pH: 7.4; 3-4 hrs) to remove residual antifreeze solution. Following this wash period, sections were subjected to an antigen retrieval step (0.05 M sodium citrate in TBS; 30 min), blocked in blocking solution [20% normal goat serum (NGS), 0.3% Triton-X, 1% H_2_O_2_ in TBS; 1 hr], and incubated at 4°C in mouse anti-AVP (PS41; 1:100, 2% NGS, 0.3% Triton-X; overnight). Sections were rinsed in TBS and incubated in a biotinylated secondary antibody solution [goat anti-mouse (1:500, BA-9200, Vector), 2% NGS, 0.3% Triton-X in TBS; 1 hr]. Tissue sections were next incubated in avidin-biotin complex (ABC Elite Kit; PK-6100, Vector; 1 hr) and visualized using diaminobenzadine (DAB peroxidase substrate kit; SK-4100, Vector; 2-3 min). Sections were mounted on gelatin-coated slides, rinsed in 50% ethanol, air-dried, and coverslipped using Permount mounting medium (SP15-100, Fisher Scientific). Slides were coded prior to image acquisition and analyzed by an experimenter blind to sex.

AVP fiber density analysis was conducted as previously described (Rood and De Vries, 2011; DiBenedictis et al., 2017). Briefly, bright-field brain section images were acquired under a 40X objective on a Keyence BZ-X700E/BZ-X710 microscope and associated BZ-H3AE software (Keyence Corporation of America), and AVP-ir fiber density was measured by manually thresholding gray-scale images of AVP-ir fibers using ImageJ (NIH, http://rsb.info.nih.gov/ij/). Brain section images were acquired under a 40X objective along the anterior-posterior axis of the VP. More specifically, AVP-ir fiber density was measured bilaterally, then averaged across hemispheres for all tissue sections within four sampling regions (Fig 5A): region 1 (+0.96 mm to +0.72 mm from bregma), region 2 (+0.60 mm to +0.36 mm from bregma), region 3 (+0.24 mm to 0.00 mm from bregma), and region 4 (−0.12 mm to −0.48 mm from bregma). Values for sections within each sampling region were then averaged in order to acquire one value per sampling region for each subject. Fiber density data are reported in number of pixels^2^, with a greater number of pixels^2^ indicating a larger area occupied by AVP-ir fibers.

### Experiment 3: Determine vasopressin 1a receptor binding density across the anterior to posterior axis of the ventral pallidum

To determine potential sex differences in V1aR binding density in the VP, V1aR binding density was measured across the anterior-posterior axis of the VP. Juvenile (35 days old) male (n = 13) and female (n = 13) rats were euthanized with CO_2_. Brains were extracted, and rapidly frozen in methylbutane cooled in dry ice and stored at −45°C. Brains were cut in 16 μm coronal sections into 8 series that were mounted directly on slides (Superfrost Plus; Fisher Scientific) and stored at −45°C until receptor autoradiography was performed. Receptor autoradiography was conducted as previously described (Lukas et al., 2010; Smith et al., 2017) using a linear V1aR antagonist [^125^I]-d(CH_2_)^5^Tyr(Me)AVP (Perkin Elmer, USA) as the tracer. Briefly, the slides from one series of tissue were thawed and dried at room temperature, followed by a short fixation in paraformaldehyde (0.1%). The slides were washed two times in 50mM Tris (pH 7.4), exposed to the tracer buffer (50 pM tracer, 50 mM Tris, 10 mM MgCl2, 0.01% BSA; 1 hr) and washed four times in Tris + 10 mM MgCl2. The slides were then briefly dipped in pure water, air-dried, exposed to Bioax MR films (Kodak), and developed after eight days. The optical density of V1aR binding was measured using ImageJ (NIH, http://rsb.info.nih.gov/ij/). Data was converted to disintegrations per minute/milligram (dpm/mg) tissue using a [^125^I] standard microscale (American Radiolabeled Chemicals Inc). Tissue background was subtracted for each measure. More specifically, V1aR binding density was measured bilaterally, then averaged across both hemispheres for all tissue sections within four sampling regions (Fig 5A): region 1 (+0.96 mm to +0.72 mm from bregma), region 2 (+0.60 mm to +0.36 mm from bregma), region 3 (+0.24 mm to 0.00 mm from bregma), and region 4 (−0.12 mm to −0.48 mm from bregma). Values for sections within each sampling region were then averaged in order to acquire one value per sampling region for each subject.

### Experiment 4: Determine whether exposure to social play enhances recruitment of V1aR-expressing ventral pallidal cells

Juvenile rats were exposed to social play in order to assess neuronal activation patterns across the anterior-posterior axis of the VP after exposure to social play, including within V1aR-expressing VP cells. Juvenile (26 days old) male and female rats were pair-housed with an age- and sex-matched cagemate, and divided into “Social Play” (males n = 6; females n = 7) and “No Social Play” (males n = 7; females n = 5) conditions. At 33 days of age, all experimental subjects were single-housed in new cages and the pair-housed cages of the experimental subjects in the “Social Play” condition were kept in order to be used as the social play testing environment. The following day, experimental subjects in the “Social Play” condition were rejoined with their previous cagemate in their original pair-housed cage for the social play test. Social play testing was otherwise conducted as described in Experiment 1 above. After the end of the 10 min session, experimental subjects were returned to their single-housed cages and remained there until transcardial perfusions. One of the experimental subjects in each pair was striped with a permanent marker 30–60 min prior to social play exposure in order to distinguish the two experimental subjects during later video analysis. Social play tests were videotaped and behavioral scoring was conducted as described in Experiment 1 above, with the following change: each video was scored twice, once for each experimental subject in the pair.

Thirty minutes after the start of the social play test, experimental subjects in the “Social Play” condition were euthanized via transcardial perfusions. This time course was chosen because stimulus-induced *c-fos* mRNA expression is at its peak 30 min after stimulation (Morgan and Curran, 1991). Experimental subjects in the “No Social Play” condition remained single-housed and undisturbed until euthanasia via transcardial perfusions. Perfusion and postfixation procedures were as described in Experiment 2, with the exception that brains were stored at −80°C.

Once mounted in the cryostat (Leica CM3050 S), brains were blocked using a razor blade in order to collect tissue containing the VP (corresponding to distances +0.96 mm to −0.24 mm from bregma; Paxinos and Watson, 2007) from the right hemisphere only (the left hemisphere was used for purposes outside the scope of this study). Brains were sliced in 30μm coronal sections into 4 series on a cryostat and each series was collected into TBS (50mM, pH: 7.4) and mounted on separate slides (Superfrost Plus; Fisher Scientific) on the same day, which were then stored at −80°C until subsequent histological processing.

The first tissue series was processed for *in situ* hybridization using an RNAScope^™^ Multiplex Fluorescent Reagent Kit v2 (323100, Advanced Cell Diagnostics) to quantify *fos* and *v1aR* mRNA expressing cells according to user manual from the supplier (Document Number 323100-USM, Advanced Cell Diagnostics). Briefly, tissue sections were washed in phosphate buffer solution (pH 7.6), dried at 60°C (30 min), then post-fixed in 4% paraformaldehyde followed by dehydration in an ethanol series. Following hydrogen peroxide incubation and target retrieval in a steamer, tissue was then treated with protease III (30 min; 322340, Advanced Cell Diagnostics) at room temperature. *fos*-C1 (403591, Advanced Cell Diagnostics) and *v1aR-C3* (402531-C3, Advanced Cell Diagnostics) probes were then hybridized in a HybEZ^™^ oven (2 hrs; Advanced Cell Diagnostics) at 40°C. After probe hybridization, tissue sections were incubated with amplifier probes (AMP1, 40°C, 30 min; AMP2, 40°C, 30 min; AMP3, 40°C, 15min). *fos* mRNA was then tagged to the fluorophore fluorescein (1:1500; NEL741E001KT, Akoya Biosciences), and *v1aR*-mRNA was tagged to the fluorophore Cy5 (1:1500; NEL745E001KT, Akoya Biosciences). Slides were then rinsed in TBS and stained with a fluorescent Nissl (1:500, 1 hr; NeuroTrace^™^; N21479, 435/455nm, Thermo Fisher Scientific). Slides were then coverslipped with Vectashield hardset antifade mounting medium with a DAPI counterstain (H-1500-10, Vector Laboratories) and stored at 4°C.

All images were acquired with a 40x objective on a Keyence BZ-X700E/BZ-X710 fluorescent microscope and associated BZ-H3AE software (Keyence Corporation of America). Images were taken at eight sampling locations across the VP in order to assess the distribution of *v1aR*+ cells, *fos*+ cells, and *v1aR*+ cells that co-expressed *fos* in the VP. Specifically, the following anterior-posterior distances were imaged (distances refer to mm from bregma; Paxinos and Watson, 2007): +0.84 mm, +0.48 mm, +0.12 mm, and −0.24 mm, and at each anterior-posterior distance a dorsal image and a ventral image were taken (Fig. 2A). Cells were counted as *v1aR*+ if they had one or more puncta, and as *fos*+ if they had five or more puncta (Fig. 2B). In addition to counting the number of *v1aR*+ cells, *fos*+ cells, and *v1aR*+ cells that co-express *fos* for each image, the percent of *v1aR*+ cells that co-expressed *fos* was quantified as [(# double-labelled cells/total number of *v1aR*+ cells)*100] and the percent of *fos*+ cells that co-expressed *v1aR* was quantified as [(# double-labelled cells/total number of *fos*-mRNA cells)*100].

**Figure 2.**
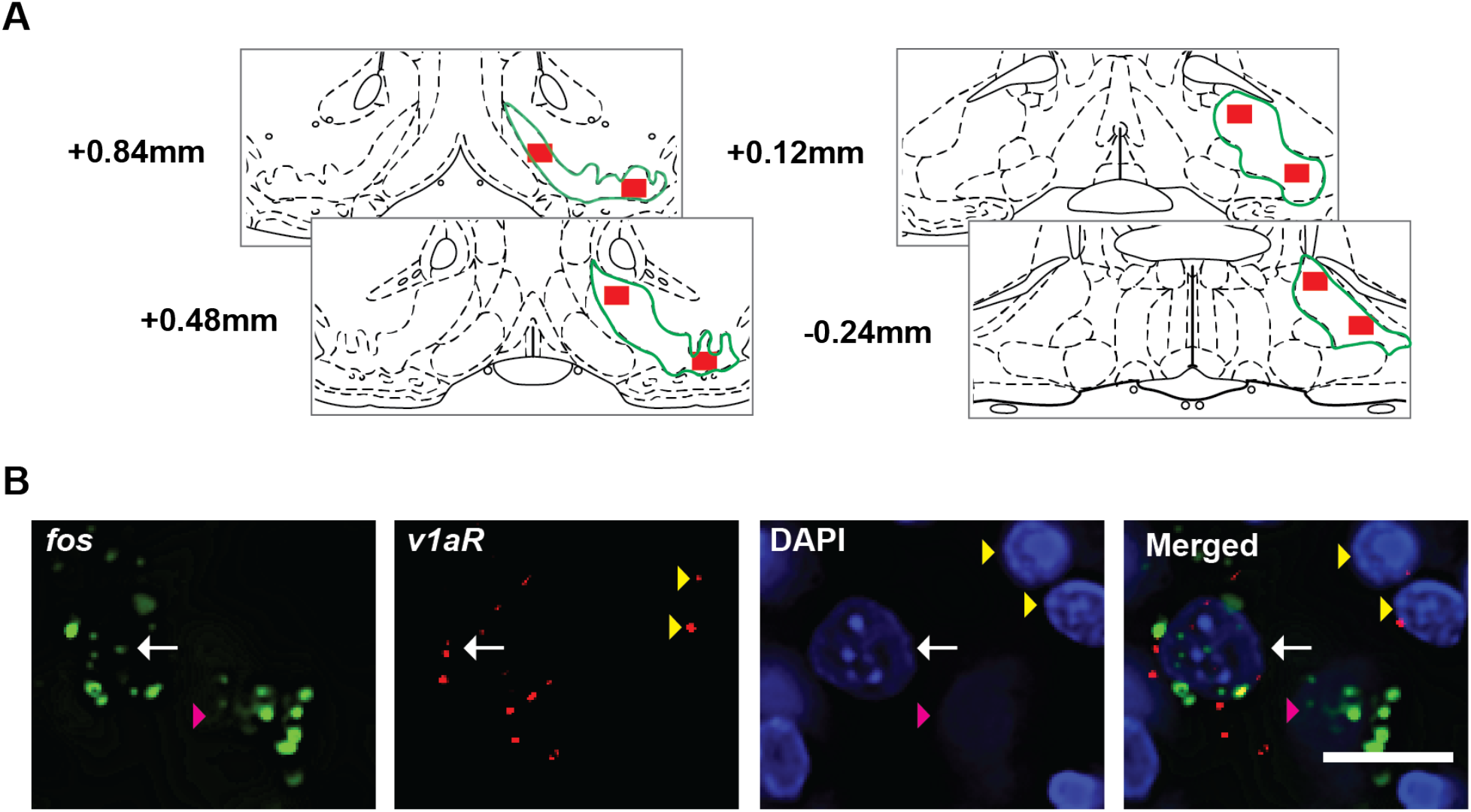
(**A**) Image sampling locations in the ventral pallidum for Experiment 4 shown on modified rat brain atlas templates (Paxinos and Watson, 2007). Ventral pallidum is outlined in green; numbers to the left of the atlas templates represent distance from Bregma; red boxes represent the sampling location where images were taken. (**B**) Example photomicrographs showing *fos* mRNA staining in green, *v1aR* mRNA staining in red, and DAPI nuclear counterstain in blue. The number of single-labelled *fos*+ cells (pink arrowheads), single-labelled *v1aR*+ cells (yellow arrowheads), and double-labelled cells (white arrow) cells were counted. Scale bar = 10 μm.

### Experiment 5: Determine the involvement of vasopressin signaling in the ventral pallidum in the regulation of social play behavior

In order to determine whether pharmacological manipulation of AVP signaling in the VP alters the expression of social play behavior, a selective V1aR antagonist or synthetic AVP was bilaterally infused into the VP prior to exposure to the social play test. Juvenile (31 days old) male and female rats underwent stereotaxic surgery for the bilateral implantation of guide cannulae targeting the VP. The stereotaxic coordinates and surgical protocols were conducted as described in Experiment 1, with the exception that some rats received s.c. injections of meloxicam (Covetrus, 049756; 2mg/kg) instead of Rimadyl immediately after surgery and once a day for an additional two days. Two days after surgery, experimental subjects (33 days old) received bilateral infusions of vehicle [artificial cerebrospinal fluid (aCSF); pH 7.4; males n = 12; females n = 15)], the specific V1aR antagonist d(CH2)5Tyr(Me2)AVP (Manning et al., 2008; 5ng/hemisphere, males n = 6; females n = 7), or synthetic AVP (Sigma Aldrich, V9879-5MG; 400pg/hemisphere, males n = 8; females n = 9) into the VP 20 min prior to exposure to the social play test as described in Experiment 1. Time of administration and concentrations of drugs for the behavioral test were based on previous studies showing that these drugs caused changes in social behaviors of juvenile rats (Veenema et al., 2012, 2013).

Social play testing, behavioral analysis, and histological verification of cannulae placements were conducted as described in Experiment 1. Only rats with bilateral (V1aR-α) or unilateral (vehicle or AVP) cannulae tracks terminating in the VP were included in the final analyses (Fig. 3). The rationale for this is that typically, bilateral infusions of an antagonist are necessary to induce a behavioral effect (Seamans et al., 1998) while unilateral infusions of an agonist can be sufficient to induce a behavioral effect (Lee et al., 1998). Furthermore, our preliminary analysis indicated that there was no main effect of unilateral versus bilateral cannulae placements on social play behaviors within the AVP treatment group.

**Figure 3.**
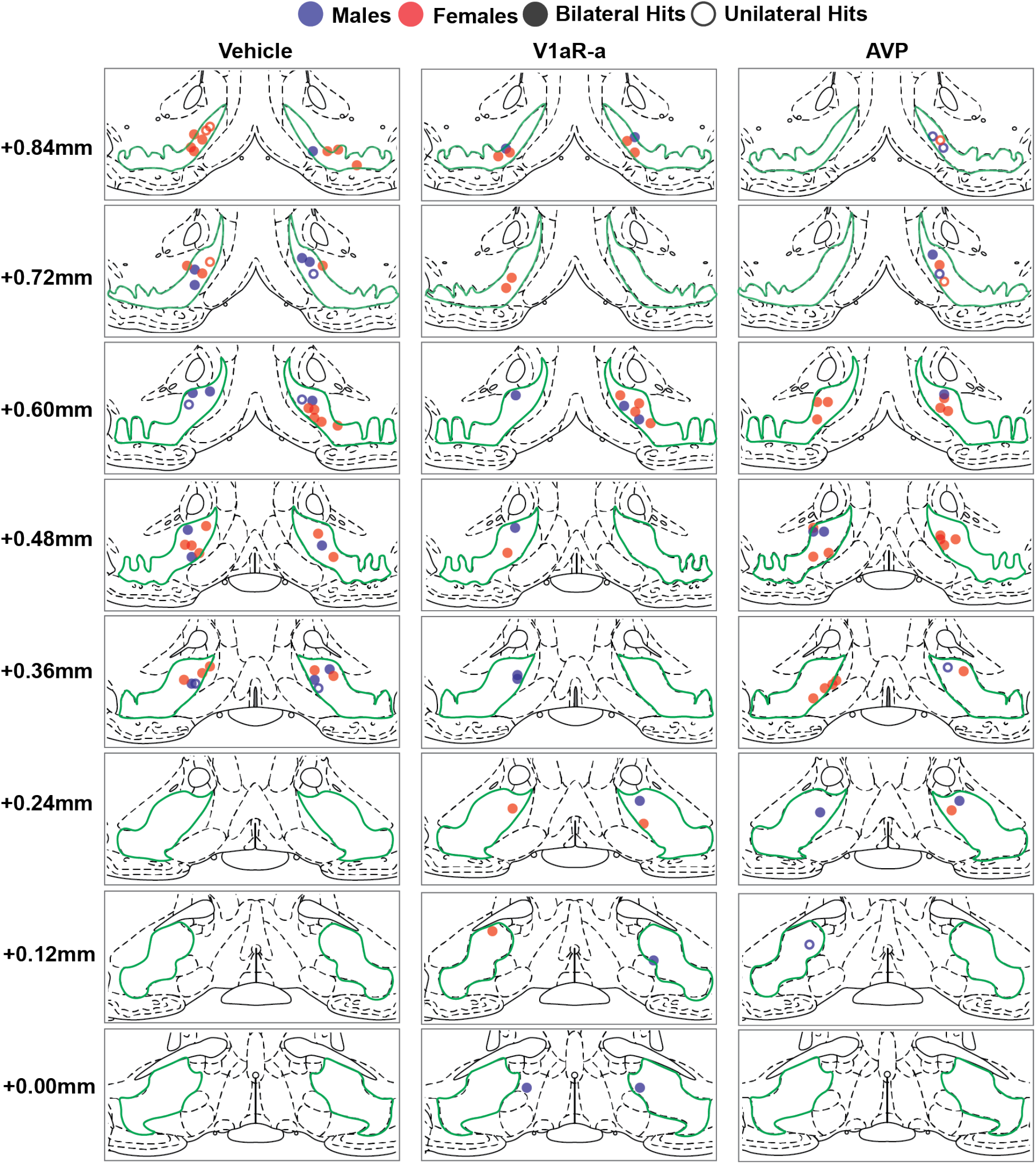
Bilateral or unilateral cannulae placements in the ventral pallidum (based on the central placement of charcoal that was injected as marker) for Experiment 5 shown on modified rat brain atlas templates (Paxinos and Watson, 2007). Incorrect contralateral cannulae placements for unilateral hits can be found in Supplementary Figure 1 for vehicle and AVP. Ventral pallidum is outlined in green; numbers to the left of the atlas templates represent distance from Bregma.

### Statistical analysis

For Experiment 1, a two-way analysis of variances (ANOVA) was used to determine the effects of drug treatment (between-subjects factor) and sex (between-subjects factor) on social play expression. For Experiments 2 and 3, mixed-effects models (REML) were used to analyze the effects of sampling region (within-subjects factor) and sex (between-subjects factor) on AVP-ir fiber density and V1aR binding density. For Experiment 4, independent samples t-tests were used to analyze the effect of sex on behaviors during the social play test, mixed-model [anterior-posterior location (within-subjects factor) x dorsal-ventral location (within-subjects factor) x sex (between-subjects factor) x social play condition (between-subjects factor)] ANOVAs were used to assess the effects of sex and social play exposure on activation of the VP, and Pearson correlations were used to determine whether activation in the VP correlated with the percent of time experimental subjects in the “Social Play” condition engaged in social play. For Experiment 5, a two-way ANOVA was used to determine the effects of drug treatment (between-subjects factor) and sex (between-subjects factor) on social play expression. When significant interactions were found in the ANOVAs and mixed-effects models, Bonferroni *post hoc* multiple comparison tests were conducted to clarify the effects. Normality was assessed by Shapiro-Wilk testing of the raw data, sphericity by Mauchly’s Test, and equality of variances by Levene’s Test. All data were analyzed using Graphpad Prism 9 or IBM SPSS 26, and statistical significance was set at *p* < 0.05. Cohen’s d (*d*) effect sizes for all t-tests, and partial eta squared (η_p_^2^) effect sizes for all ANOVAs and mixed effects models were manually computed when significant main effects or interactions were found.

## Results

### Experiment 1: Ventral pallidal infusions of the GABA_A_ receptor agonist muscimol decreased the expression of social play behavior in both sexes

Subjects that received bilateral ventral pallidal infusions of muscimol exhibited lower levels of social play behaviors when compared to subjects that received the vehicle, as reflected by a significant main effect of drug treatment on social play duration (*F*_(1,20)_ = 19.78, *p* < 0.001, η_p_^2^ = 0.32; Fig. 4A), the number of nape attacks (*F*_(1,20)_ = 24.08, *p* < 0.001, η_p_^2^ = 0.35; Fig. 4B), and the number of pins (*F*_(1,20)_ = 10.67, *p* < 0.05, η_p_^2^ = 0.25; Fig. 4C). Muscimol did not alter the number of supine poses (*p* > 0.05; Suppl. Fig. 2A). In addition, the duration of other social behaviors during the test, specifically social investigation and allogrooming, was similar between vehicle- and muscimol-treated subjects (*p* > 0.05, both; Fig. 4D & E). Non-social cage exploration was significantly higher in muscimol-treated subjects compared to vehicle-treated subjects (*F*_(1,20)_ = 8.33, *p* < 0.05, η_p_^2^ = 0.23; Fig. 4F). There were no main effects of, or interactions with, sex in the expression of any social or non-social behaviors (*p* > 0.05, all).

**Figure 4.**
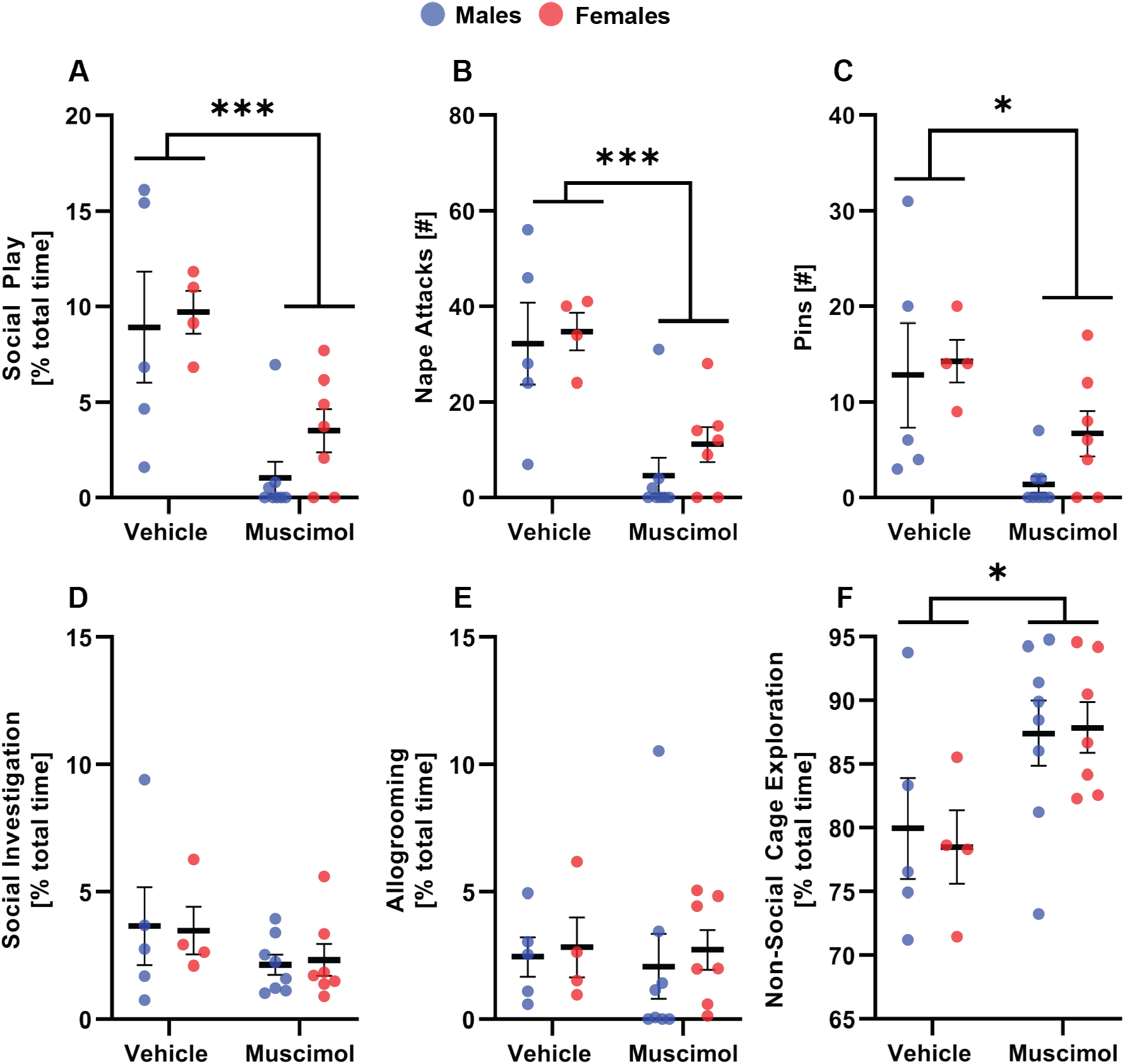
Bilateral infusion of muscimol into the VP decreased the expression of social play behaviors in male and female rats. Bilateral ventral pallidal infusions of the GABA_A_ receptor agonist muscimol decreased the duration of social play (**A**), the number of nape attacks (**B**), and the number of pins (**C**) in male and female rats compared to their vehicle-treated counterparts. Muscimol increased the duration of non-social cage exploration in male and female rats compared to vehicle-treated counterparts **(F**), but did not alter the duration of social investigation (**D**) or allogrooming (**E**). Black bars indicate mean ± SEM; \**p* < 0.05, \*\*\**p* < 0.001, two-way ANOVA.

### Experiment 2: Vasopressin-ir fibers are denser in males than in females across the analyzed regions of the ventral pallidum

AVP-ir fiber density was significantly different between males and females across the analyzed regions of the VP. There was a significant main effect of sex (*F*_(1,6)_ = 6.01, *p* < 0.05, η_p_^2^ = 0.50; Fig. 5C), with males showing denser AVP-ir fibers than females. AVP-ir fiber density was consistent across the anterior to posterior sampling regions of the VP, as reflected by no main effect of, or interaction with, sampling location on AVP-ir fiber density (*p* > 0.05, both).

**Figure 5.**
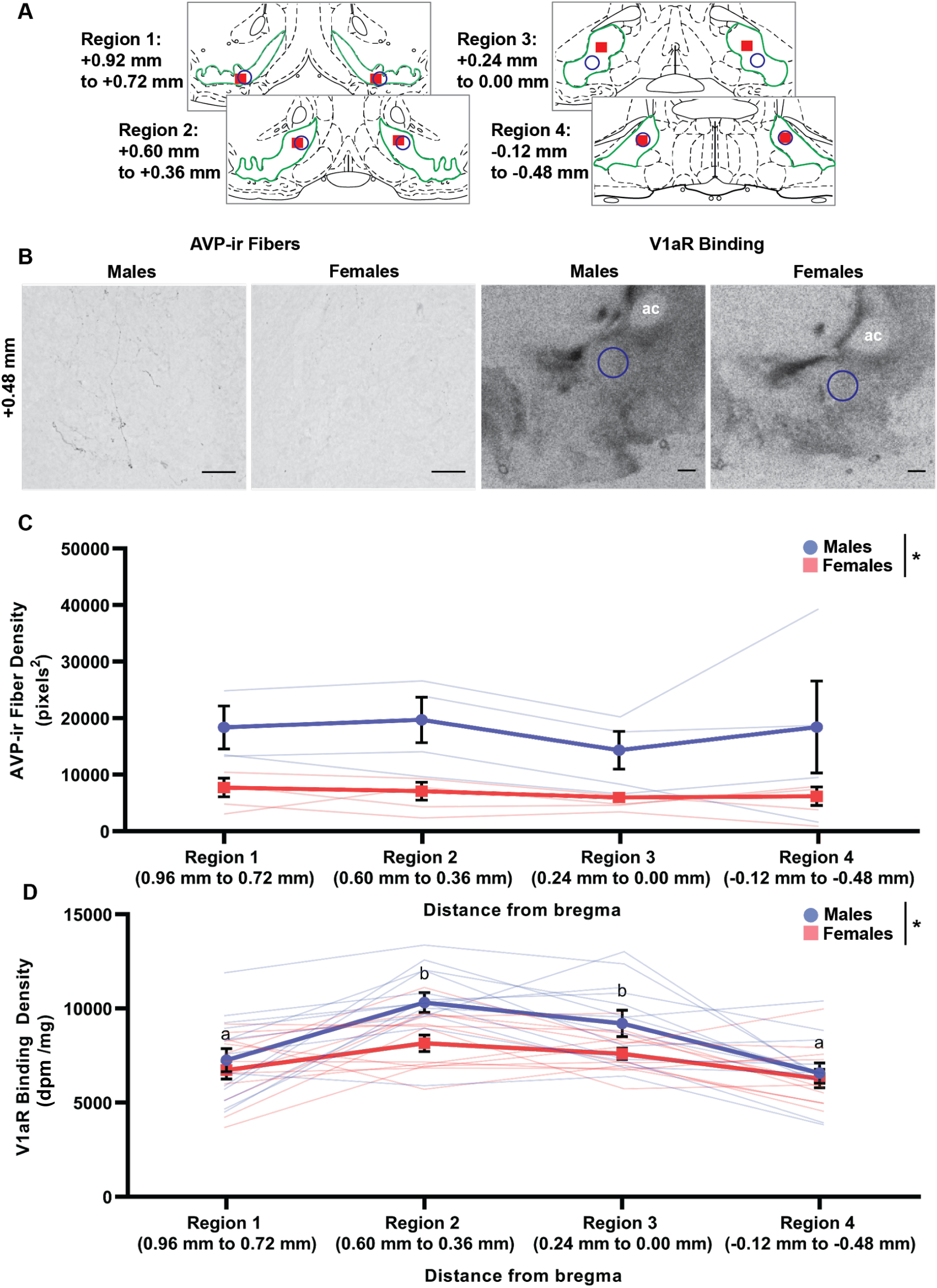
AVP-ir fiber and V1aR binding densities across the analyzed regions of the VP in male and female rats. Image sampling locations in the ventral pallidum for Experiments 2 and 3 shown on modified rat brain atlas templates (Paxinos and Watson, 2007); ventral pallidum is outlined in green; red-filled boxes represent approximate sampling locations where images were taken for the fiber analyses for each region; blue circles represent approximate sampling locations where measurements were taken for receptor binding analyses for each region; numbers to the left of the atlas templates represent distances from Bregma (Paxinos and Watson, 2007). Representative images from the right hemisphere in sampling region 2 showing AVP-ir fibers and V1aR binding in the VP of juvenile male and female rats (**B**). Males had denser AVP-ir fibers than females across the anterior to posterior sampling regions of the VP (**C**). Males had denser V1aR binding than females across the anterior to posterior sampling regions of the VP, and V1aR binding density changed across the VP, with denser binding in regions 2 and 3 compared to regions 1 and 4 in both sexes (**D**). Data shown as mean ± SEM; \**p* < 0.05, mixed-effects model; regions with different letters are significantly different from one another, Bonferroni *post hoc* comparisons; scale bars = 15 μm in **B** and 500 μm in **C**; ac, anterior commissure.

### Experiment 3: Vasopressin 1a receptor binding is denser in males than in females and changes across the analyzed regions of the ventral pallidum

V1aR binding density differed between males and females, and across the analyzed regions of the VP. In detail, there was a significant main effect of sex (*F*_(1,24)_ = 5.98, *p* < 0.05, η_p_^2^ = 0.20; Fig. 5D), with males showing denser V1aR binding in the VP than females. There was also a significant main effect of sampling location (*F*_(2.35, 52.40)_ = 15.85, *p* < 0.001, η_p_^2^ = 0.42; Fig. 5D) on V1aR binding density, but no sex X sampling location interaction (*p* > 0.05). *Post hoc* tests indicated that V1aR binding density was similar between regions 2 and 3 (*p* > 0.05), and between regions 1 and 4 (*p* > 0.05). However, regions 2 and 3 showed denser V1aR binding than regions 1 and 4 (*p* < 0.05, all; Fig. 5D).

### Experiment 4: Exposure to social play enhanced recruitment of V1aR-expressing ventral pallidal cells in males but not in females

Males and females showed similar duration of social play (*p* > 0.05; Fig. 6A), number of pins (*p* > 0.05; Fig. 6C), number of supine poses (*p* > 0.05; Suppl. Fig. 2B), duration of social investigation (*p* > 0.05; Fig. 6D) and duration of non-social cage exploration (p > 0.05; Fig. 6F). However, males showed a greater number of nape attacks (*t*_(14)_ = 2.23, *p* < 0.05, *d* = 1.11; Fig. 6B) and a longer duration of allogrooming (*t*_(14)_ = 2.51, *p* < 0.05, *d* = 1.26; Fig. 6E) than females.

**Figure 6.**
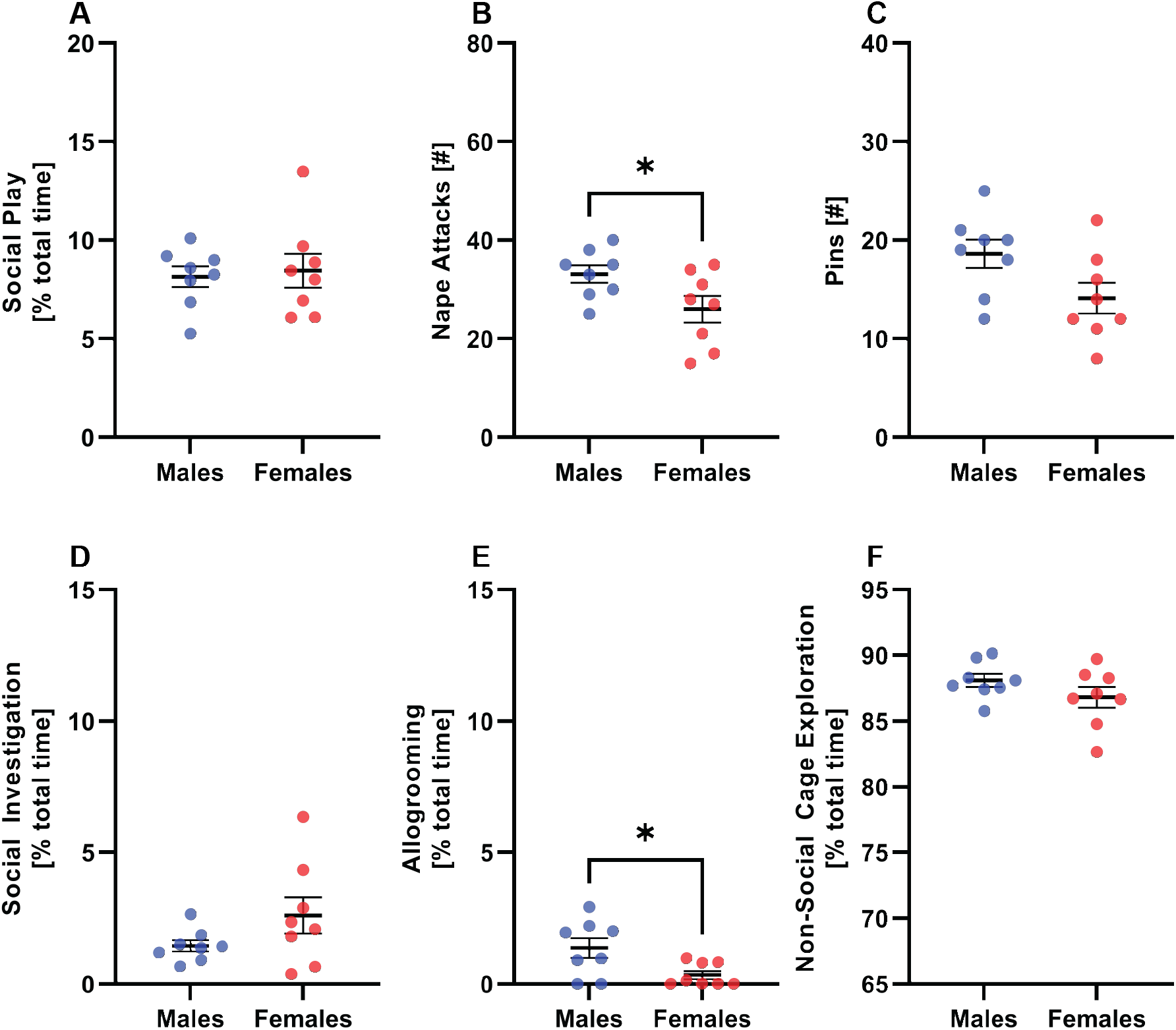
Expression of social behaviors during the social play test in sex- and age-matched cagemates. There was no sex difference in the duration of social play (**A**), the number of pins (**C**), the duration of social investigation (**D**), or the duration of non-social cage exploration (**F**). Males showed higher number of nape attacks (**B**) and duration of allogrooming (**E**) compared to females. Black bars indicate mean ± SEM; \**p* < 0.05, independent samples t-test.

There was an overall sex difference in the number of *v1aR*-expressing cells in the VP. Females had a greater number of *v1aR*+ cells in the VP compared to males, as reflected by a significant main effect of sex (*F*_(1,21)_ = 7.69, *p* < 0.05, η_p_^2^ = 0.27; Fig. 7A, B). There were no significant interactions with sex for any other factors (*p* > 0.05 all). There was also a significant main effect of dorsal-ventral location (*F*_(1,21)_ = 36.37, *p* < 0.0001, η_p_^2^ = 0.63; Fig. 7A, B) and a significant dorsal-ventral location X anterior-posterior location interaction (*F*_(1,21)_ = 7.62, *p* < 0.05, η_p_^2^ = 0.27; Fig. 7A), but no main effect of anterior-posterior location (*p* > 0.05). *Post hoc* tests confirmed that there was a greater number of *v1aR*+ cells in the dorsal location compared to the ventral location at each anterior-posterior location (*p* > 0.05, all; Fig 7A, B). Further, *post hoc* tests indicated that the number of *v1aR*+ cells was consistent across the anterior-posterior sampling locations in the VP; there were no significant differences in the number of *v1aR*+ cells between anterior-posterior locations for either the dorsal location or the ventral location (*p* > 0.05 all). There were no main effects of, or interactions with, social play condition on the number of *v1aR*+ cells (*p* > 0.05 all).

**Figure 7.**
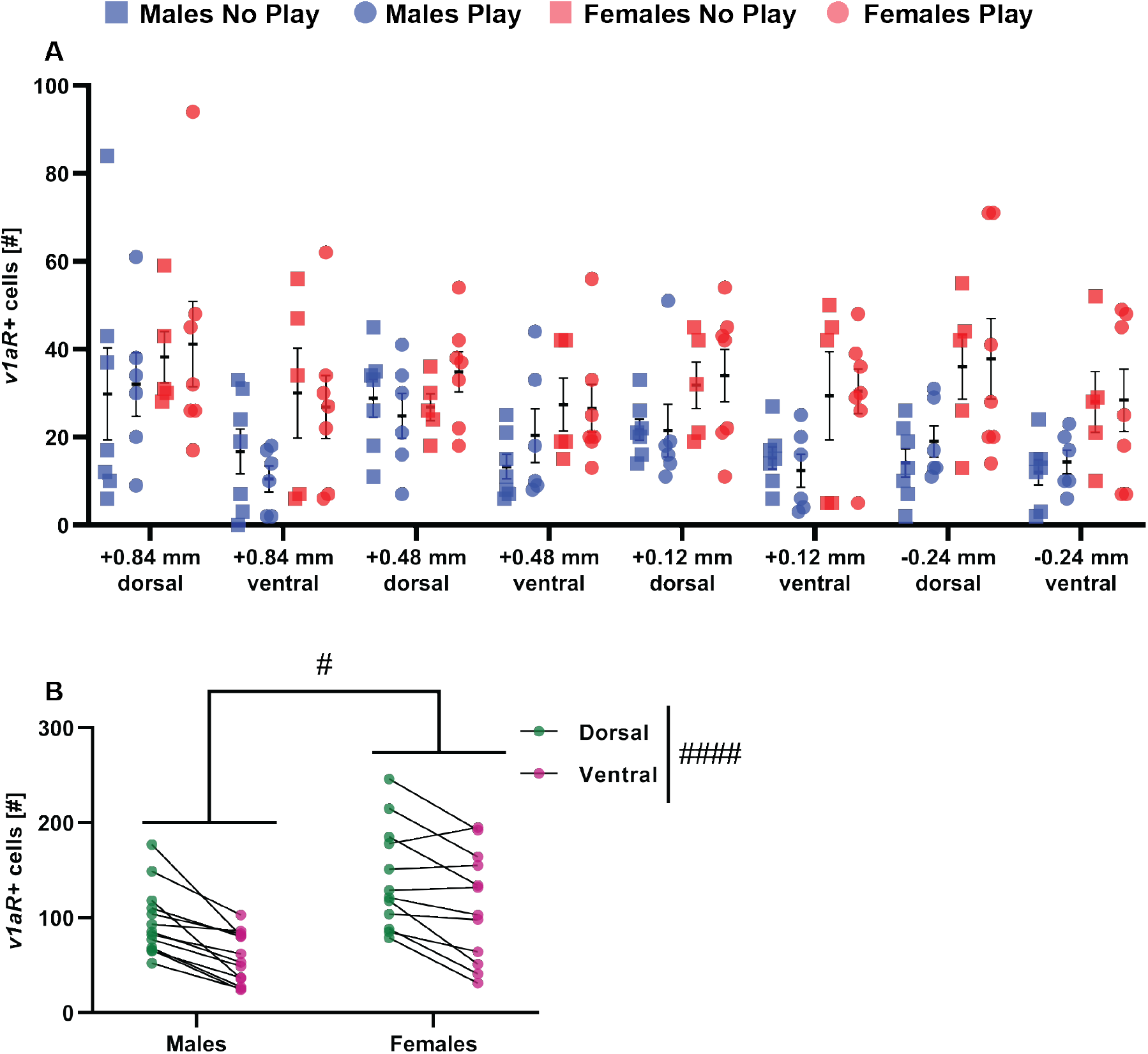
Distribution of V1aR-expressing cells across the analyzed regions of the VP in male and female rats. Females showed a greater number of *v1aR*+ cells than males across the anterior to posterior sampling regions of the VP (**A & B**). In addition, there was a greater number of *v1aR*+ cells in the dorsal locations compared to the ventral locations in both sexes (**A & B**). Data is shown across anterior-posterior and dorsal-ventral locations in **A**, and collapsed across anterior-posterior location and social play condition to highlight significant main effects in **B**. Black bars indicate mean ± SEM; #*p* < 0.05, #### *p* < 0.0001, mixed-model ANOVA main effects.

Exposure to social play recruited VP cells in a sex-specific manner. There were no main effects of sex or social play condition, but a significant sex X social play condition interaction for the total number of *fos*+ cells in the VP (*F*_(1,21)_ = 4.87, *p* < 0.05, η_p_^2^ = 0.19; Fig. 8A, B). *Post hoc* tests indicated that males in the “Social Play” condition had significantly more *fos*+ cells than males in the “No Social Play” condition and females in the “Social Play” condition (*p* < 0.05, both; Fig. 8A, B). The number of *fos*+ cells was similar in females from the “Social Play” and “No Social Play” conditions (*p* > 0.05; Fig. 8A, B), and between males and females in the “No Social Play” condition (*p* > 0.05; Fig. 8A, B). There were no main effects of, or interactions with, anterior-posterior location or dorsal-ventral location on *fos* expression (*p* > 0.05 all; Fig. 8A). The duration of social play did not significantly correlate with the number of *fos*+ cells when the data was collapsed by sex (*p* > 0.05). However, a negative correlation between social play duration and the number of *fos*+ cells approached significance in males (*r*_(6)_ = 0.80, *p* = 0.054; Fig. 8C), and a positive correlation between social play duration and the number of *fos*+ cells approached significance in females (*r*_(7)_ = 0.70, *p* = 0.079; Fig. 8D).

**Figure 8.**
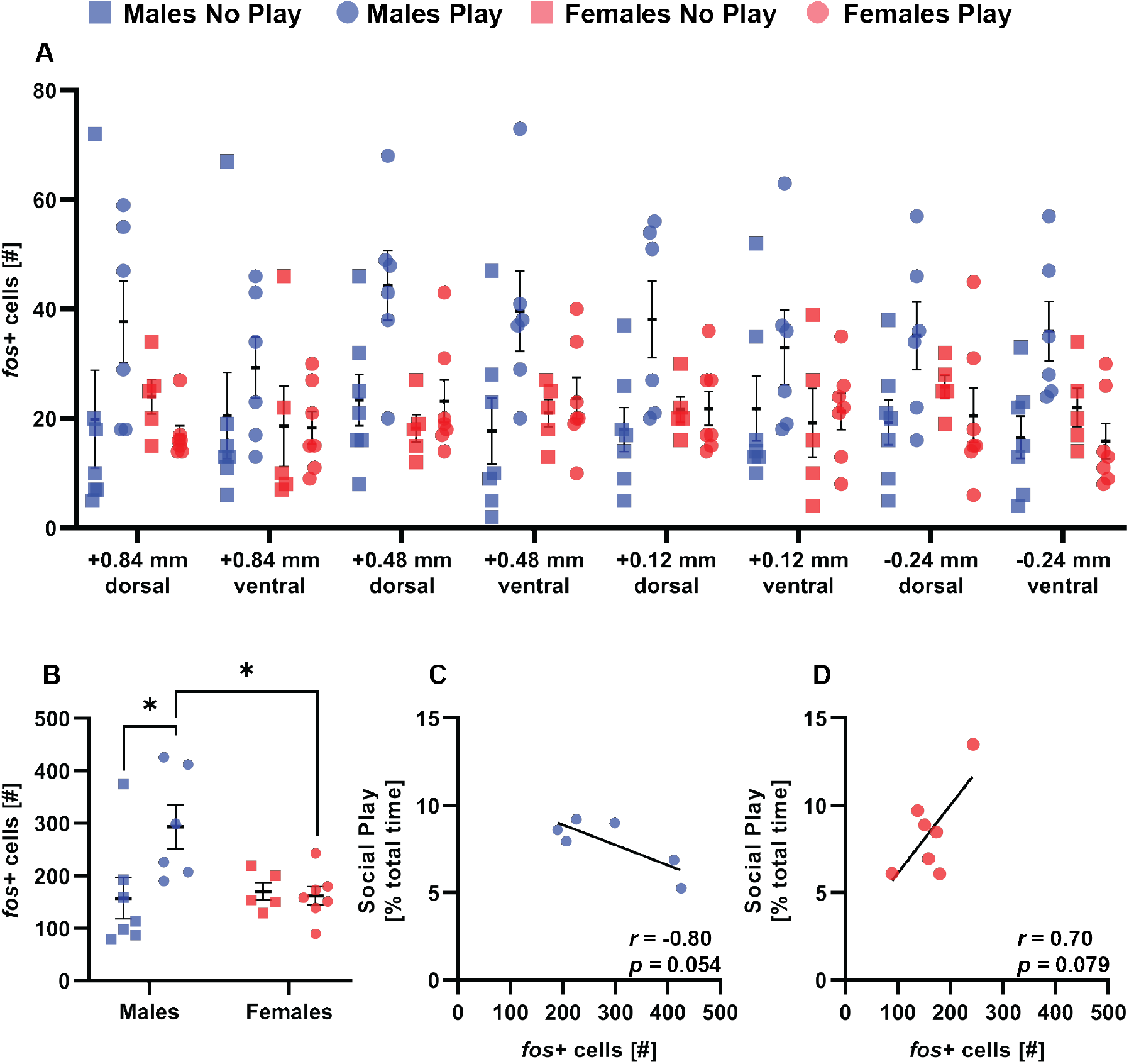
VP cells were recruited in a sex-specific manner by exposure to social play. Exposure to social play enhanced recruitment of VP cells (**A** & **B**) in males, but not in females. Time spent engaging in social play did not significantly correlate with the number of *fos*+ cells in males (**C**) or females (**D**). The number of *fos*+ cells was similar across sampling regions in all groups (**A**), thus data were collapsed across sampling regions to highlight sex and play condition effects in **B**. Black bars indicate mean ± SEM; \**p* < 0.05, Bonferroni *post hoc* comparisons.

Exposure to social play also recruited VP *v1ar*+ cells in a sex-specific manner. There was a significant main effect of social play condition (*F*_(1,21)_ = 9.43, *p* < 0.05, η_p_^2^ = 0.31; Fig. 9A, B) and a significant sex X social play condition interaction (*F*_(1,21)_ = 26.84, *p* < 0.0001, η_p_^2^ = 0.56; Fig. 9A, B), but no main effet of sex (*p* > 0.05). Subjects in the “Social Play” condition had a greater proportion of *v1aR*+ cells that co-expressed *fos* compared to subjects in the “No Social Play” condition, however *post hoc* tests indicated that this effect was driven by the males. Specifically, males in the “Social Play” condition had a greater proportion of *v1aR*+ cells that coexpressed *fos* than males in the “No Social Play” condition (*p* < 0.05; Fig. 9A, B), but the proportion of *v1aR*+ cells that co-expressed *fos* was similar between females in the “Social Play” and “No Social Play” conditions (*p* > 0.05; Fig. 9A, B). *Post hoc* tests also indicated that males in the “Social Play” and “No Social Play” conditions exhibited a greater proportion of *v1aR*+ cells that co-expressed *fos* than their female counterparts (*p* < 0.001, and *p* < 0.05 respectively; Fig. 9A, B). The duration of social play did not significantly correlate with the proportion of *v1aR*+ cells that co-expressed *fos* when data were collapsed or separated by sex (*p* > 0.05 all; Fig. 9C & D).

**Figure 9.**
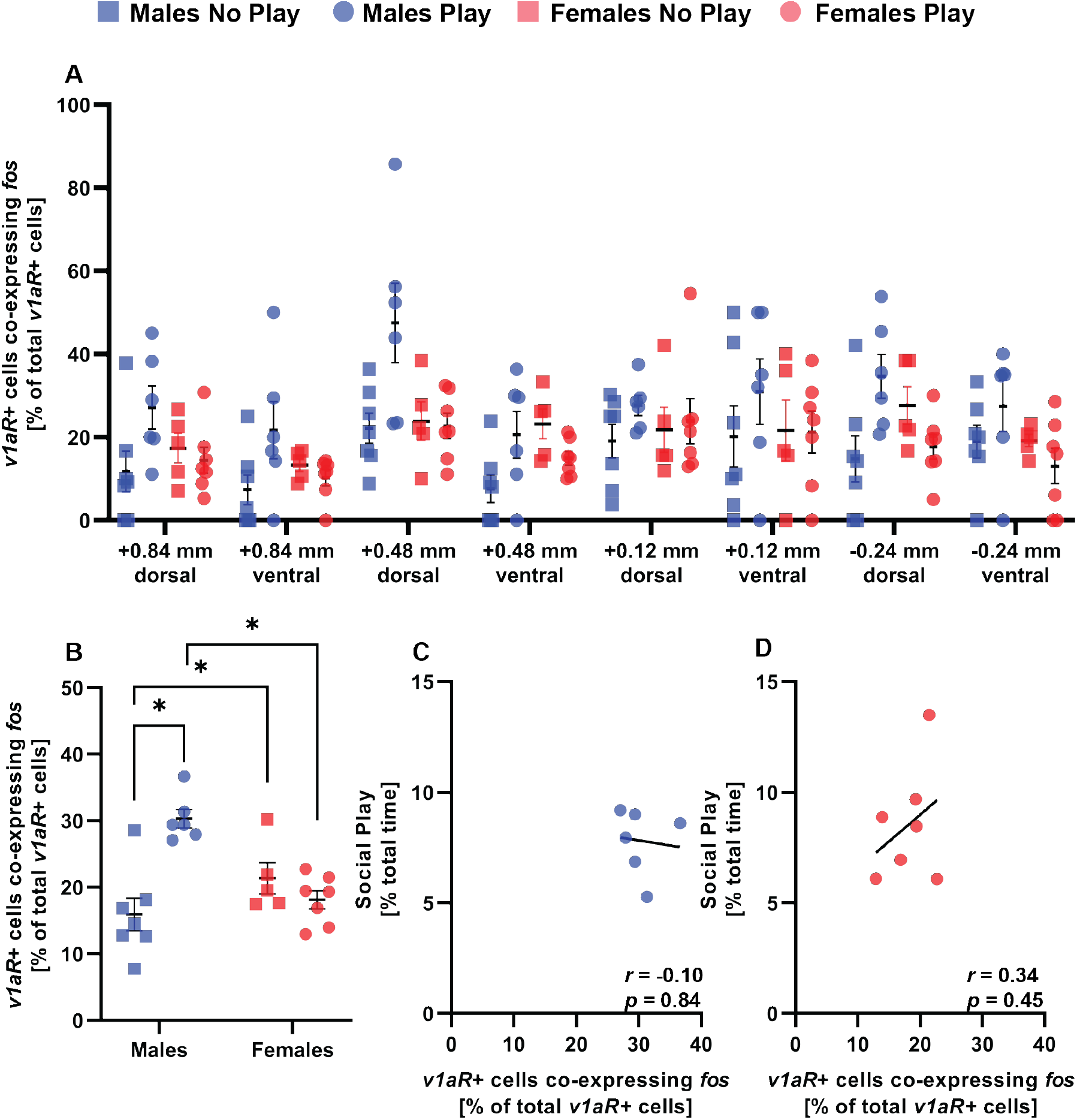
Exposure to social play recruited V1aR-expressing VP cells in a sex-specific way. Males exposed to social play showed a greater proportion of *v1aR*+ cells that co-expressed *fos* compared to males that were not exposed to social play and females that were exposed to social play (**B**). Time spent engaging in social play did not significantly correlate with the proportion of *v1aR*+ cells that co-expressed *fos* in males (**C**) or females (**D**). Data is shown across anterior-posterior and dorsal-ventral locations in **A**, and collapsed across sampling locations to highlight sex and social play condition effects in **B**. Black bars indicate mean ± SEM; *\*p* < 0.05, Bonferroni *post hoc* comparisons.

The proportion of *v1aR*+ cells that co-expressed *fos* varied across anterior-posterior and dorsal-ventral locations, but these effects were independent of sex and social play condition. There was a significant main effect of anterior-posterior location (*F*_(3,63)_ = 5.32, *p* < 0.05, η_p_^2^ = 0.20; Fig. 9A). *Post hoc* tests indicated that there was a greater proportion of *v1aR*+ cells that coexpressed *fos* at bregma distances 0.48 mm and 0.12 mm compared to bregma distance 0.84 mm (*p* < 0.05); no other paired comparisons reached significance (*p* > 0.05, all). There was also a main effect of dorsal-ventral location (*F*_(1,63)_ = 5.13, *p* < 0.05, η_p_^2^ = 0.07; Fig. 9A), with the dorsal locations exhibiting a greater proportion of *v1aR*+ cells that co-expressed *fos* than the ventral locations (*p* < 0.05; Fig. 9A). There were no significant interactions between anterior-posterior location and dorsal-ventral location, nor did these factors interact with sex or social play condition (*p* > 0.05, all).

Exposure to social play did not alter the proportion of *fos*+ cells that co-expressed *v1aR*. There was a significant main effect of sex (*F*_(1,21)_ = 17.50, *p* < 0.001, η_p_^2^ = 0.45; Fig. 10B), with females exhibiting a greater proportion of *fos*+ cells that co-expressed *v1aR* than males. There was also a significant main effect of dorsal-ventral location (*F*_(1,63)_ = 17.54, *p* < 0.001, η_p_^2^ = 0.22; Fig. 10A), with the dorsal locations exhibiting a greater proportion of *fos*+ cells that coexpressed *v1aR* than the ventral locations. There were no main effects of social play condition or anterior-posterior location, nor any significant interactions between any factors (*p* > 0.05 all). The duration of social play did not significantly correlate with the proportion of *fos*+ cells that co-expressed *v1aR* when collapsed or separated by sex (*p* > 0.05 all; Fig. 10C & D).

**Figure 10.**
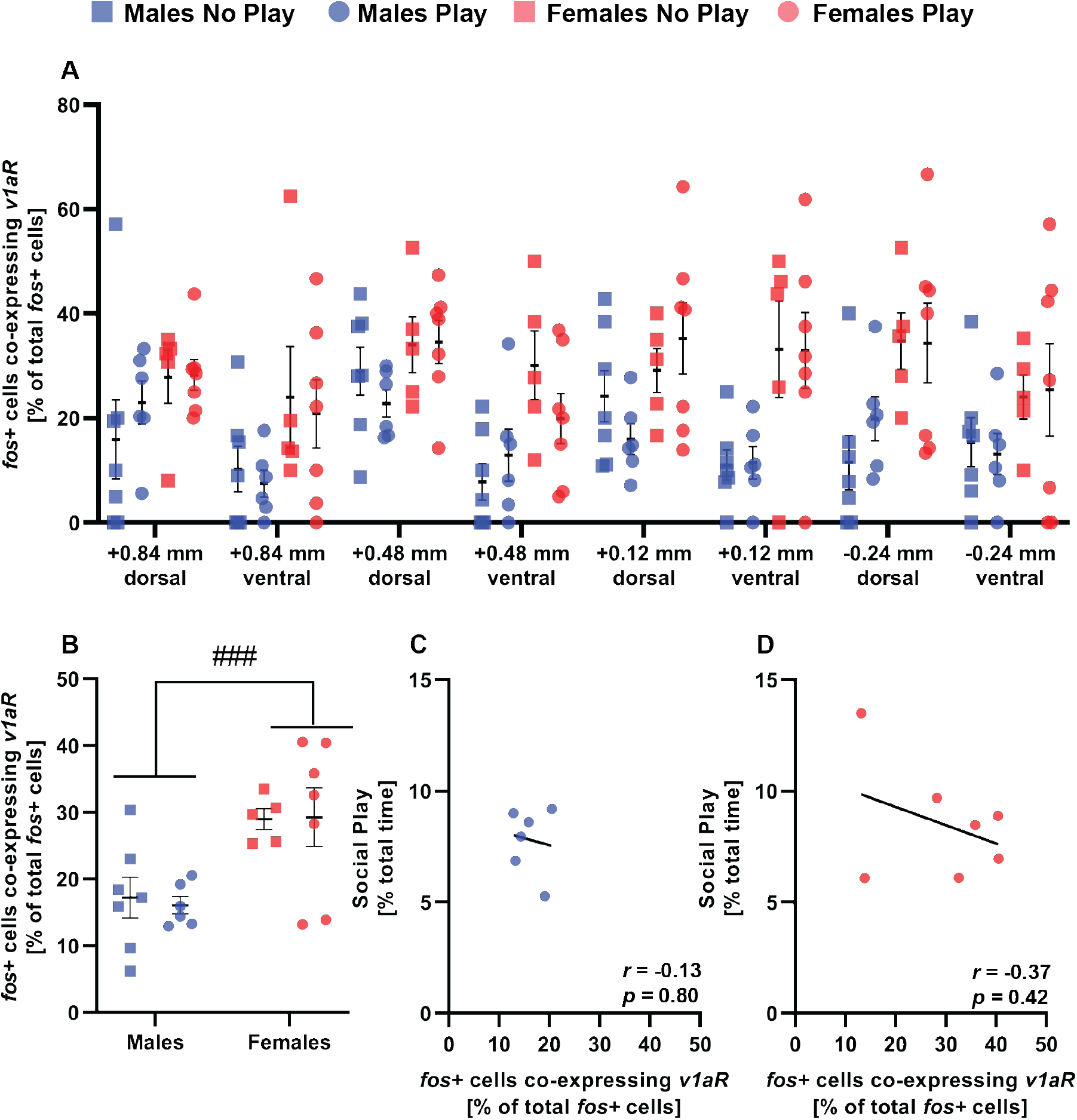
Exposure to social play did not alter the proportion of *fos*+ cells that coexpressed *v1aR*. Females showed a greater proportion of *fos*+ cells that co-expressed *v1aR* than males (**B**). Time spent engaging in social play did not significantly correlate with the proportion of *fos*+ cells that co-expressed *v1aR* in males (**C**) or females (**D**). Data is shown across anterior-posterior and dorsal-ventral locations in **A**, and collapsed across sampling locations to highlight sex and social play condition effects in **B**. Black bars indicate mean ± SEM; ###*p* < 0.001, mixed-model ANOVA main effect

### Experiment 5: V1aR antagonist treatment in the ventral pallidum altered the expression of social play behavior in sex-specific ways

Pharmacological blockade of V1aR’s in the VP, via bilateral infusions of the V1aR antagonist, altered social play in sex-specific ways. There was a significant sex X drug interaction for the duration of social play (*F*_(2,51)_ = 9.35, *p* < 0.001, η_p_^2^ = 0.21; Fig. 11A), the number of nape attacks (*F*_(2,51)_ = 16.65, *p* < 0.0001, η_p_^2^ = 0.26; Fig. 11B), and the number of pins (*F*_(2,51)_ = 13.65, *p* < 0.0001, η_p_^2^ = 0.24; Fig. 11C). *Post hoc* tests indicated that V1aR antagonist treatment altered social play behaviors in sex-specific ways while synthetic AVP treatment had no effect on the expression of social play behaviors. Specifically, V1aR antagonist-treated males, compared to vehicle- and AVP-treated males, showed longer social play duration (*p* < 0.05, both; Fig. 11A), more nape attacks (*p* < 0.0001, both; Fig. 11B), and more pins (*p* < 0.001, both; Fig. 11C). In contrast, V1aR antagonist-treated females, compared to vehicle- and AVP-treated females, showed shorter social play duration (*p* < 0.05, both; Fig. 11A). In addition, V1aR antagonist-treated females, compared to vehicle-treated females, showed fewer nape attacks (*p* < 0.05; Fig. 11B), and fewer pins (*p* < 0.05; Fig. 11C). V1aR antagonist-treated males showed longer social play duration and more nape attacks and pins than V1aR antagonist-treated females (*p* < 0.001, all; Fig. 11A & B), creating a sex difference that was not observed in the vehicle group (p > 0.05, all).

**Figure 11.**
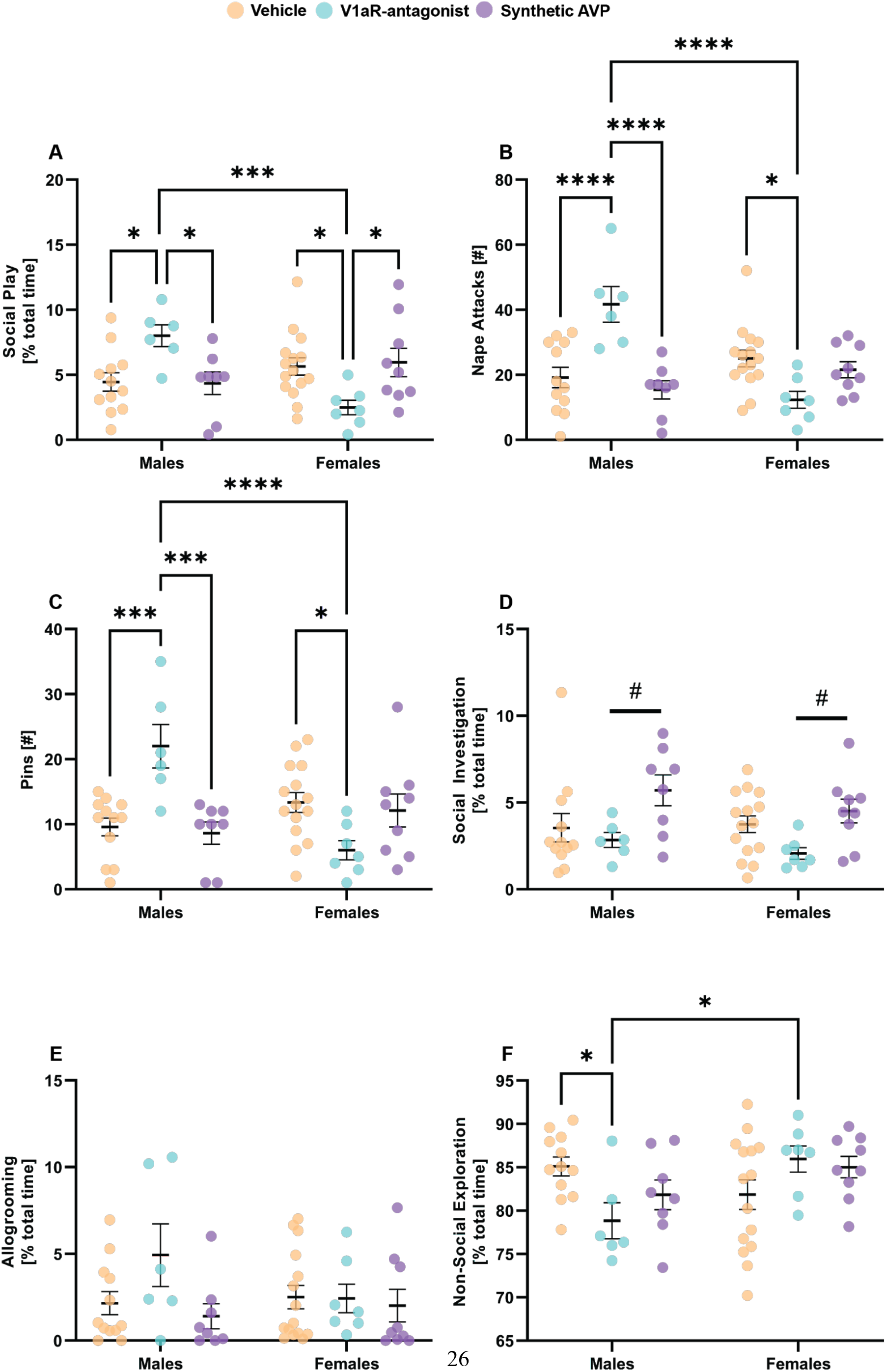
Pharmacological manipulation of AVP signaling in the VP altered social play behaviors in sex-specific ways. Bilateral ventral pallidal infusions of the specific V1aR antagonist increased social play duration (**A**), the number of nape attacks (**B**), and the number of pins (**C**) in male rats but reduced these parameters in juvenile female rats when compared to vehicle-treated counterparts. Synthetic AVP-treated subjects showed a higher duration of social investigation when compared to their V1aR antagonist-treated counterparts (**D**). V1aR antagonist or synthetic AVP treatment did not alter the duration of allogrooming (**E**). V1aR antagonist treatment reduced the duration of non-social cage exploration in males only when compared to their vehicle-treated counterparts (**F**). Black bars indicate mean ± SEM; \**p* < 0.05, \*\*\**p* < 0.001, \*\*\*\**p* < 0.0001, Bonferroni *post hoc* paired comparisons; #*p* < 0.05 Bonferroni *post hoc* comparison collapsed across sex.

Synthetic AVP did not alter the expression of social play behaviors as indicated by a similar duration of social play (*p* > 0.05; Fig. 11A), a similar number of nape attacks (*p* > 0.05; Fig. 11B), and a similar number of pins (*p* > 0.05; Fig. 11C) between AVP- and vehicle-treated subjects of both sexes.

There were no main effects of, or interactions with, sex or drug treatment on the number of supine poses (*p* > 0.05; Suppl. Fig. 2C) or the duration of allogrooming (*p* > 0.05; Fig. 11E). There was a significant main effect of drug treatment on the duration of social investigation (*F*_(2,51)_ = 6.01, *p* < 0.05, η_p_^2^ = 0.02; Fig. 11D), but no main effect of, or interaction with, sex (*p* > 0.05, both; Fig. 11D). *Post hoc* tests indicated that the duration of social investigation in AVP-or V1aR antagonist-treated subjects was similar to that of vehicle-treated subjects (*p* > 0.05; Fig. 11D), but that synthetic AVP-treated subjects showed a longer duration of social investigation than V1aR antagonist-treated subjects (*p* < 0.05; Fig. 11D). There was a significant sex X drug interaction for non-social cage exploration (*F*_(2,51)_ = 5.24, *p* < 0.05, η_p_^2^ = 0.14; Fig. 11F). *Post hoc* tests indicated that V1aR antagonist-treated males spent less time engaged in non-social cage exploration compared to vehicle-treated males and V1aR antagonist-treated females (*p* < 0.05; Fig. 11F); no other comparisons reached significance for non-social cage exploration (*p* > 0.05, all).

## Discussion

Our study indicates, for the first time, the involvement of the VP in the regulation of social play behavior. We showed that social play behavior was reduced in both male and female juvenile rats following inactivation of the VP using the GABA_A_ receptor agonist muscimol. However, only males showed an increased activation of the VP, as reflected by more *fos*+ cells, in response to social play exposure. We further found that males showed denser AVP-ir fibers and V1aR binding in the VP than females, but that females expressed more *v1aR*+ cells in the VP than males. Additionally, males, but not females, showed enhanced recruitment of *v1aR-* expressing VP cells in response to social play exposure. Finally, we found that blockade of AVP signaling in the VP using a specific V1aR antagonist increased social play duration in males, but decreased social play duration in females. Together, these findings provide the first evidence that the VP modulates social play behavior in juvenile rats and does so in a sex-specific manner through AVP signaling.

### Activation of the VP is critical for the typical expression of social play behavior

We found that temporary inactivation of the VP, through activation of GABA_A_ receptors, significantly reduced social play behaviors in both male and female juvenile rats. This reduction in social play behavior was not due to a general decrease in locomotion, because muscimol-treated male and female rats showed an increase in the duration of non-social cage exploration as compared to their vehicle-treated counterparts. We also found that the effects of muscimol were specific to social play, because the duration of social investigation and allogrooming did not change following muscimol administration. These results demonstrate the importance of the VP for the expression of social play behavior.

We found that there was a sex difference in the activation of the VP in response to social play exposure. Here, males exposed to social play showed a greater number of *fos*+ cells in the VP compared to males that were not exposed to social play. In contrast, females exposed and not exposed to social play showed a similar number of *fos*+ cells in the VP. However, a previous study found that juvenile male rats exposed to social play showed similar c-Fos activation in the VP compared to those that were not exposed to social play (females were not used; van Kerkhof et al., 2014). This inconsistency could be due to some key differences in the methodology between the previous study (van Kerkhof et al., 2014) and the current study. In the earlier study, previously-paired subjects were exposed to social play in a test cage (after four consecutive days of 30 min habituation sessions to the test cage), while the previously-paired subjects in the current study were exposed to social play in their home cage. This is a key methodological difference, because it has been shown that the neural regulation of social play is dependent on the social context of the testing environment (Bredewold et al., 2014). Furthermore, the previous study counted cells that were positive for the protein c-Fos, while we counted cells that were positive for the mRNA transcript *fos*. In addition, the previous study counted c-Fos+ cells only in one subregion of the VP (corresponding to −0.40 mm from bregma; Paxinos and Watson, 1998), and this subregion was more posterior to any of the multiple sampling regions (from +0.84 mm to −0.24 mm from bregma; Paxinos and Watson, 2007) examined for *fos+ cells* in the current study. These differences in methodology may account for the different results reported between the two studies. We also found that in males, the duration of social play and the number of *fos*+ cells in the VP showed a negative correlation approaching significance, while in females there was a positive correlation approaching significance between the two parameters. These findings suggest that social play is regulated by a decrease in VP activation in males, but an increase in VP activation in females. Taken together, our pharmacological and *in situ* hybridization data demonstrate that temporary inactivation of the VP decreased social play behaviors in both sexes, while exposure to social play recruited VP cells in a sex-specific manner.

The VP is part of the SDMN, a network of brain regions regulating rewarding social behaviors (O’Connell and Hofmann, 2011). Several other brain structures in the SDMN have also been implicated in the regulation of social play behavior in rats (Meaney et al., 1981; Schneider and Koch, 2005; Taylor et al., 2012; van Kerkhof et al., 2014; Veenema et al., 2013; van Kerkhof et al., 2013; Bredewold et al., 2014, 2015; Paul et al., 2014; Manduca et al., 2016; Argue et al., 2017; Reppucci et al., 2018; Zhao et al., 2020), suggesting that multiple brain regions form a circuit to modulate the expression of social play behavior. The VP receives major input from the nucleus accumbens (NAc; Williams et al., 1977; Nauta et al., 1978; Zahm and Heimer, 1990; Heimer et al., 1991), which is also part of the SDMN (O’Connell and Hofmann, 2011) and a well-known key node in the mesocorticolimbic reward system (Kelley and Berridge, 2002; Berridge and Kringelbach, 2015). The VP and NAc modulate various rewarding social behaviors such as pair-bonding (Pitkow et al., 2001; Liu and Wang, 2003; Lim and Young, 2004), maternal behavior (Numan et al., 2005; Olazábal and Young, 2006), sociosexual motivation (Beny-Shefer et al., 2017; DiBenedictis et al., 2020), and social play behavior (van Kerkhof et al., 2013, 2014; Manduca et al., 2016); current study). Given that both the NAc and VP are independently involved in the regulation of rewarding social behaviors, we hypothesize that the NAc to VP pathway mediates these social behaviors. In future experiments, it would be of interest to test whether and how the NAc to VP pathway regulates social play and other socially rewarding behaviors.

### Sex differences in the AVP system in the VP of juvenile rats

We report in the current study that juvenile male rats had denser AVP-ir fibers in the VP than juvenile female rats. This is in line with similar sex differences in AVP-ir fiber density in the VP found in adult rats and prairie voles, with males showing denser AVP-ir fibers than females (Lim et al., 2004a; DiBenedictis et al., 2020). The sex difference in AVP-ir fiber density in the VP may be due to sex differences in AVP input to the VP. In support, in adult rats, AVP input to the VP originates in the BNST and medial amygdala (MeA; DiBenedictis et al., 2020), with adult males having more AVP-ir cells (van Leeuwen et al., 1985; Miller et al., 1989; Wang et al., 1993; DiBenedictis et al., 2017), and a greater percentage of VP-projecting AVP-ir cells (DiBenedictis et al., 2020) than adult females. It would be of interest to determine whether the BNST and MeA also send AVP projections to the VP in juvenile rats, and whether there are sex differences in the strength of these projections. Furthermore, to understand its functional implications, it will be important to determine whether the observed sex difference in AVP-ir fiber density in the VP corresponds with a sex difference in AVP release into the VP.

We also report in the current study that juvenile males show denser V1aR binding in the VP than juvenile females. Our findings are in contrast to an earlier study, which reported no sex differences in V1aR binding in the VP of juvenile rats (Smith et al., 2017). However, this earlier study only examined V1aR binding at a single location (corresponding to 1.56 mm from bregma, Paxinos and Watson, 2007) which was more anterior to the multiple sampling regions analyzed in the current study (0.96 mm to −0.48 mm from bregma, Paxinos and Watson, 2007). Prior studies in adult rodents reported no sex differences in V1aR binding density in rats or Taiwan meadow and forest voles (Chappell et al., 2016; Smith et al., 2017). However, it should be noted that it is unclear from the studies in adult rodents whether V1aR binding density varies between the sexes across the anterior to posterior axis of the VP because only one location within the VP was analyzed. Even so, these findings may suggest a potential role of the sex difference in V1aR binding in the VP in the juvenile stage only, although this remains to be tested.

We found that there was a sex difference in the number of *v1aR*-expressing cells in the VP, with females exhibiting a greater number of VP *v1aR*-expressing cells than males. This sex difference in the number of *v1aR*-expressing cells is opposite of the sex difference in V1aR binding density (with males showing denser V1aR binding than females). It is important to point out that V1aR binding density and the number of *v1aR-*positive cells are two different types of measurements. V1aR receptor binding measures V1aR receptor affinity (for review see: Kuhar et al., 1986) irrespective of the number of cells expressing the V1aR, while measuring the number of cells that express *v1aR* mRNA doesn’t take into account V1aR affinity. Additionally, V1aR receptor binding could represent V1aRs on presynaptic terminals of VP afferents or on postsynaptic membranes of VP cells (Pittman et al., 1998; Frey and Albin, 2001; Oz et al., 2001; Bailey et al., 2006), while *v1aR* expression was determined by proximity to nuclei and thus likely limited to VP cells (for review see: Ben-Yishay and Shav-Tal, 2019; Stavast and Erkeland, 2019). Taken together, the observed sex differences in AVP-ir fiber density, V1aR binding density, and the number of *v1aR-*positive cells in the VP of juvenile rats suggest that males, compared to females, receive more AVP projections and exhibit higher levels of V1aR affinity, but have fewer cells that express the V1aR. Future studies are needed to determine how these structural sex differences in the AVP system in the VP correspond with functional sex differences in AVP signaling in the VP.

### AVP signaling in the VP is involved in the sex-specific regulation of social play behavior

We found that V1aR antagonist treatment altered social play in a sex-specific manner. V1aR antagonist-treated males showed higher social play behaviors (i.e. increased duration, higher number of nape attacks and pins) while V1aR antagonist-treated females showed lower social play behaviors (i.e. decreased duration, fewer number of nape attacks and pins) when compared to their vehicle-treated counterparts. In contrast, V1aR antagonist treatment did not affect the expression of other social behaviors in the behavior test compared to vehicle treatment, demonstrating that V1aR blockade specifically altered the expression of social play behaviors. This sex-specific regulation of social play via AVP signaling was previously observed in the LS (Bredewold et al., 2014; Veenema et al., 2013). Here, blockade of AVP signaling via the V1aR antagonist in the LS also increased social play behaviors in males, but decreased social play behaviors in females (Bredewold et al., 2014; Veneema et al., 2013). These findings together demonstrate a unifying role of the V1aR in the LS and VP in social play behavior, such that activation of the V1aR exerts an inhibitory effect on social play in juvenile males, but a facilitatory effect on social play in juvenile females.

Synthetic AVP treatment in the VP did not alter the expression of social play behaviors in either sex. It might be the case that a higher dose of synthetic AVP is required to alter social play behavior, or that V1aRs were occupied at max capacity by endogenous AVP, leaving no receptors available to bind to additional AVP molecules. Yet, we found that intra-VP application of synthetic AVP increased the duration of social investigation when compared with V1aR antagonist-treated counterparts. Application of synthetic AVP in the LS and olfactory bulb improves social recognition memory as reflected by increased investigation of a novel as opposed to a familiar conspecific in juvenile and adult rats (Dluzen et al., 1998; Veenema et al., 2012) or decreased investigation of a familiar conspecific in adult rats (Dantzer et al., 1988). The effects of synthetic AVP on social investigation in the current study and these previous studies, suggest the potential involvement of intra-VP AVP signaling in regulating social recognition in juvenile rats, although this remains to be tested.

We also found that exposure to social play induced a sex difference in the activation of *v1aR*-expressing cells in the VP. In detail, males exposed to social play had more *v1aR*-expressing cells that co-expressed *fos* compared to males not exposed to social play. In contrast, females showed no change in recruitment of *v1aR*-expressing cells after exposure to social play. These findings, together with the V1aR antagonist findings, further emphasize the complex sexspecific involvement of the V1aR in the VP in the regulation of social play behavior. In males, the increased social play behaviors by VP-V1aR blockade and the enhanced recruitment of V1aR-expressing cells in the VP upon social play exposure suggest a potential inhibitory effect of V1aR activation on social play behavior. In females, despite an absence of a change in the recruitment of V1aR-expressing cells in the VP upon exposure to social play, the decreased social play behaviors by VP-V1aR blockade suggest a potential facilitatory effect of V1aR activation on social play behavior. Future studies incorporating *in vitro* and/or *in vivo* electrophysiological recordings are needed to determine if and how v1aR-expressing cells in the VP of males and females respond differently to application of the V1aR antagonist.

Although AVP signaling in the VP has been implicated in the regulation of adult-specific social behaviors, such as pair-bonding (Pitkow et al., 2001; Lim and Young, 2004) and sociosexual motivation (DiBenedictis et al., 2020), this is the first study to demonstrate its involvement in the regulation of a juvenile-typical social behavior. This suggests the versatile involvement of the AVP system in the VP in social behaviors across the lifespan. Furthermore, the current study and the study by DiBenedictis et al. (2020) demonstrate that AVP signaling in the VP modulates social behaviors in sex-specific ways in juveniles and adults. Specifically, in the current study, blocking V1aR’s in the VP induced a sex difference that was not seen under vehicle conditions. We propose that the structural sex differences in the AVP system in the VP contribute to preventing a sex difference in social play behavior. This would support the Dual-Function Hypothesis of neural sex differences in which it is suggested that sex differences in the brain may serve two functions: to induce overt sex differences in behavior or to prevent overt sex differences in behavior (De Vries, 2004). How these structural sex differences in the AVP system in the VP are functionally linked to the sex-specific regulation of social play behavior remains to be tested.

### Future directions: Determine the source of AVP input to the VP

Although AVP-ir cells in the BNST and MeA have been shown to project to the VP in adult male and female rats (DiBenedictis et al., 2020), it is unknown whether these two brain regions send AVP projections to the VP in juvenile rats. Previous studies have shown that the BNST (Paul et al., 2014; van Kerkhof et al., 2014) and MeA (Argue et al., 2017) are involved in the regulation of social play. For example, juvenile males, but not females, show a negative correlation between AVP mRNA expression in the BNST and social play behavior (Paul et al., 2014), suggesting the potential of an inhibitory role for BNST AVP in the expression of social play behavior in males. In contrast, juvenile male and female rats showed greater recruitment of the BNST (van Kerkhof et al., 2014; Reppucci et al., 2018) and MeA (Reppucci et al., 2018) in response to social play. However it is unclear whether this activation was within AVP cells or other cell populations. To address these outstanding questions, future studies should determine whether BNST and MeA AVP cells project to the VP, and whether exposure to social play enhances recruitment of VP-projecting AVP cells in these two brain regions. Together with the current and previous findings, these studies will provide critical insights into a circuit, modulated by AVP signaling, that underlies the regulation of social play behavior in juvenile male and female rats.

## Acknowledgements

This work was supported by the National Institutes of Health (R01MH102456 to Alexa H. Veenema) and the National Science Foundation (IOS 1735934 to Alexa H. Veenema; DGE-1848730 to Jessica Lee). We would like to thank Dr. Caroline J.W. Smith, Dr. Brett T. DiBenedictis, Catherine Washington, and Abigal Barrett for technical assistance. We would also like to thank Dr. Maurice Manning and Dr. Hal Gainer for supplying the V1aR antagonist and AVP antibody, respectively.

## Supplementary Material

**Supplementary Figure 1.**
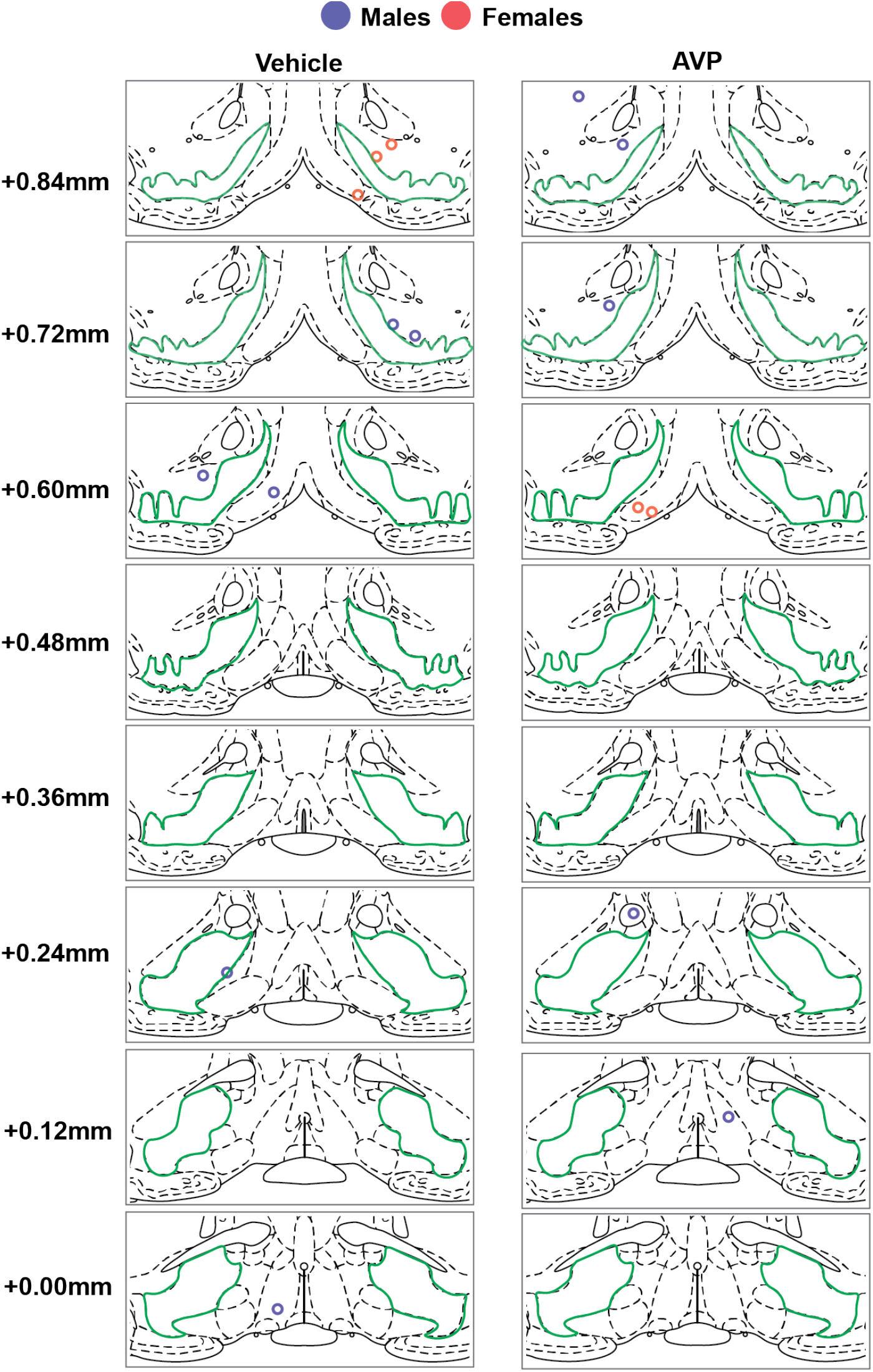
Incorrect contralateral cannulae placements for unilateral hits in the ventral pallidum (based on the central placement of charcoal that was injected as marker) for Experiment 5 shown on modified rat brain atlas templates (Paxinos and Watson, 2007). Ventral pallidum is outlined in green; numbers to the left of the atlas templates represent distance from Bregma.

**Supplementary Figure 2.**
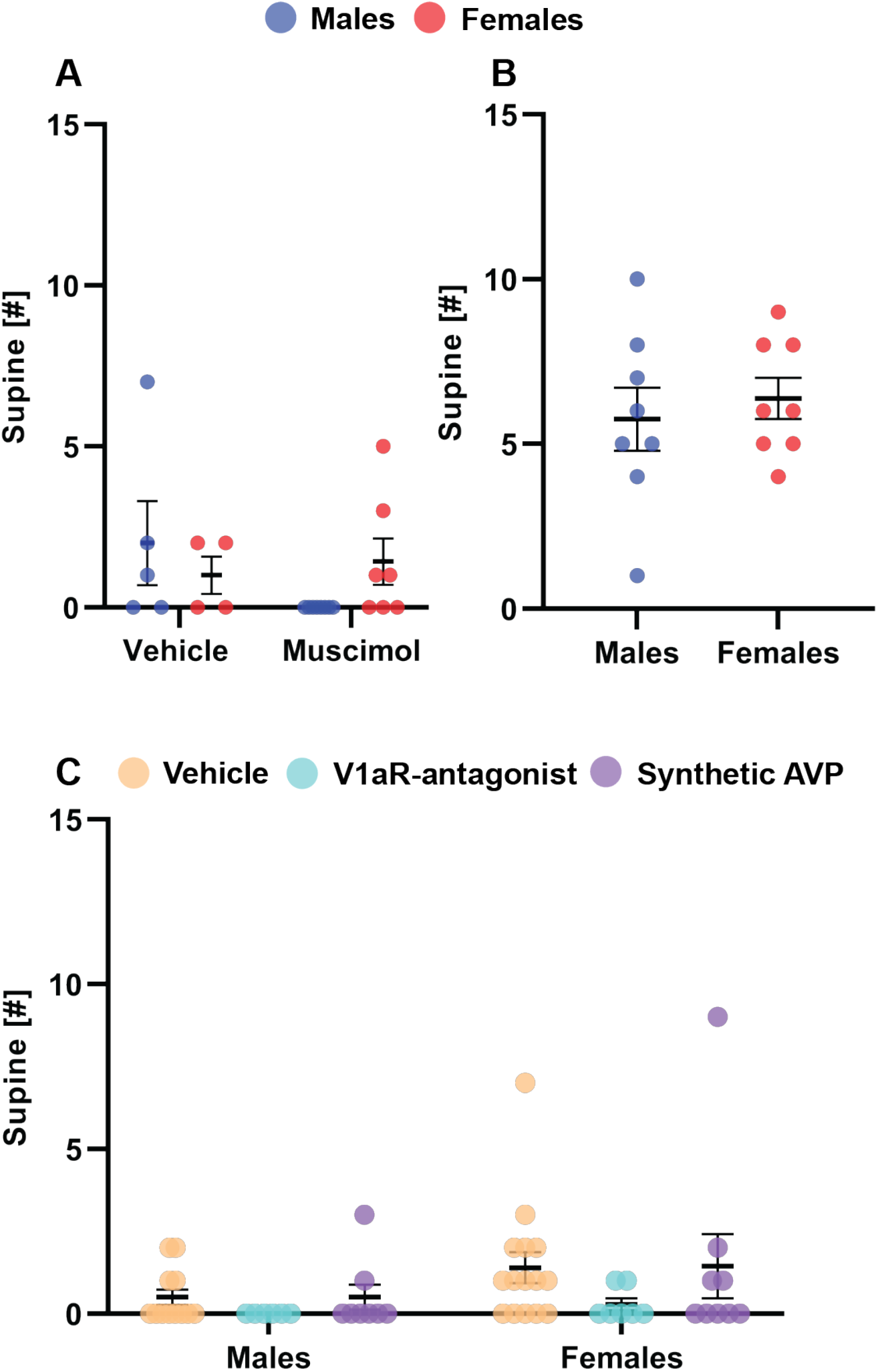
The number of supine poses from Experiment 1 (**A**), Experiment 4 (**B**), and Experiment 5 (**C**). There were no sex or drug treatment effects found in the number of supine poses for any experiment.

## Notes

### Competing Interest Statement

The authors have declared no competing interest.

## References

Argue KJ, VanRyzin JW, Falvo DJ, Whitaker AR, Yu SJ, McCarthy MM (2017) Activation of both CB1 and CB2 endocannabinoid receptors is critical for masculinization of the developing medial amygdala and juvenile social play behavior. Eneuro 4.

Bailey TW, Jin Y-H, Doyle MW, Smith SM, Andresen MC (2006) Vasopressin inhibits glutamate release via two distinct modes in the brainstem. J Neurosci 26:6131–6142.

Bekoff M (1974) Social Play and Play-Soliciting by Infant Canids. Am Zool 14:323–340.

Ben-Barak Y, Russell JT, Whitnall MH, Ozato K, Gainer H (1985) Neurophysin in the hypothalamo-neurohypophysial system. I. Production and characterization of monoclonal antibodies. J Neurosci 5:81–97.

Ben-Yishay R, Shav-Tal Y (2019) The dynamic lifecycle of mRNA in the nucleus. Curr Opin Cell Biol 58:69–75.

Beny-Shefer Y, Zilkha N, Lavi-Avnon Y, Bezalel N, Rogachev I, Brandis A, Dayan M, Kimchi T (2017) Nucleus accumbens dopamine signaling regulates sexual preference for females in male mice. Cell Rep 21:3079–3088.

Berridge KC, Kringelbach ML (2015) Pleasure systems in the brain. Neuron 86:646–664.

Bredewold R, Schiavo JK, van der Hart M, Verreij M, Veenema AH (2015) Dynamic changes in extracellular release of GABA and glutamate in the lateral septum during social play behavior in juvenile rats: Implications for sex-specific regulation of social play behavior. Neuroscience 307:117–127.

Bredewold R, Smith CJW, Dumais KM, Veenema AH (2014) Sex-specific modulation of juvenile social play behavior by vasopressin and oxytocin depends on social context. Front Behav Neurosci 8:216.

Chappell AR, Freeman SM, Lin YK, LaPrairie JL, Inoue K, Young LJ, Hayes LD (2016) Distributions of oxytocin and vasopressin 1a receptors in the Taiwan vole and their role in social monogamy. J Zool (1987) 299:106–115.

Dantzer R, Koob GF, Bluthé RM, Le Moal M (1988) Septal vasopressin modulates social memory in male rats. Brain Res 457:143–147.

De Vries GJ (2004) Minireview: Sex differences in adult and developing brains: compensation, compensation, compensation. Endocrinology 145:1063–1068.

De Vries GJ, Buijs RM (1983) The origin of the vasopressinergic and oxytocinergic innervation of the rat brain with special reference to the lateral septum. Brain Res 273:307–317.

DiBenedictis BT, Cheung HK, Nussbaum ER, Veenema AH (2020) Involvement of ventral pallidal vasopressin in the sex-specific regulation of sociosexual motivation in rats. Psychoneuroendocrinology 111:104462.

DiBenedictis BT, Nussbaum ER, Cheung HK, Veenema AH (2017) Quantitative mapping reveals age and sex differences in vasopressin, but not oxytocin, immunoreactivity in the rat social behavior neural network. J Comp Neurol 525:2549–2570.

Dluzen DE, Muraoka S, Engelmann M, Landgraf R (1998) The effects of infusion of arginine vasopressin, oxytocin, or their antagonists into the olfactory bulb upon social recognition responses in male rats. Peptides 19:999–1005.

Frey KA, Albin RL (2001) Receptor binding techniques. Curr Protoc Neurosci Chapter 1:Unit1.4.

Heimer L, Zahm DS, Churchill L, Kalivas PW, Wohltmann C (1991) Specificity in the projection patterns of accumbal core and shell in the rat. Neuroscience 41:89–125.

Kelley AE, Berridge KC (2002) The neuroscience of natural rewards: relevance to addictive drugs. J Neurosci 22:3306–3311.

Kuhar MJ, De Souza EB, Unnerstall JR (1986) Neurotransmitter receptor mapping by autoradiography and other methods. Annu Rev Neurosci 9:27–59.

Lee MD, Aloyo VJ, Fluharty SJ, Simansky KJ (1998) Infusion of the serotonin1B (5-HT1B) agonist CP-93,129 into the parabrachial nucleus potently and selectively reduces food intake in rats. Psychopharmacology 136:304–307.

Lim MM, Murphy AZ, Young LJ (2004a) Ventral striatopallidal oxytocin and vasopressin V1a receptors in the monogamous prairie vole (Microtus ochrogaster). J Comp Neurol 468:555–570.

Lim MM, Wang Z, Olazábal DE, Ren X, Terwilliger EF, Young LJ (2004b) Enhanced partner preference in a promiscuous species by manipulating the expression of a single gene. Nature 429:754–757.

Lim MM, Young LJ (2004) Vasopressin-dependent neural circuits underlying pair bond formation in the monogamous prairie vole. Neuroscience 125:35–45.

Liu Y, Wang ZX (2003) Nucleus accumbens oxytocin and dopamine interact to regulate pair bond formation in female prairie voles. Neuroscience 121:537–544.

Lukas M, Bredewold R, Neumann ID, Veenema AH (2010) Maternal separation interferes with developmental changes in brain vasopressin and oxytocin receptor binding in male rats. Neuropharmacology 58:78–87.

Lukas M, Wöhr M (2015) Endogenous vasopressin, innate anxiety, and the emission of prosocial 50-kHz ultrasonic vocalizations during social play behavior in juvenile rats. Psychoneuroendocrinology 56:35–44.

Manduca A, Servadio M, Damsteegt R, Campolongo P, Vanderschuren LJ, Trezza V (2016) Dopaminergic neurotransmission in the nucleus accumbens modulates social play behavior in rats. Neuropsychopharmacology 41:2215–2223.

Meaney MJ, Dodge AM, Beatty WW (1981) Sex-dependent effects of amygdaloid lesions on the social play of prepubertal rats. Physiol Behav 26:467–472.

Miller MA, Vician L, Clifton DK, Dorsa DM (1989) Sex differences in vasopressin neurons in the bed nucleus of the stria terminalis by in situ hybridization. Peptides 10:615–619.

Morgan JI, Curran T (1991) Stimulus-transcription coupling in the nervous system: involvement of the inducible proto-oncogenes fos and jun. Annu Rev Neurosci 14:421–451.

Nauta WJ, Smith GP, Faull RL, Domesick VB (1978) Efferent connections and nigral afferents of the nucleus accumbens septi in the rat. Neuroscience 3:385–401.

Nijhof SL, Vinkers CH, van Geelen SM, Duijff SN, Achterberg EJM, van der Net J, Veltkamp RC, Grootenhuis MA, van de Putte EM, Hillegers MHJ, van der Brug AW, Wierenga CJ, Benders MJNL, Engels RCME, van der Ent CK, Vanderschuren LJMJ, Lesscher HMB (2018) Healthy play, better coping: The importance of play for the development of children in health and disease. Neurosci Biobehav Rev 95:421–429.

Numan M, Numan MJ, Schwarz JM, Neuner CM, Flood TF, Smith CD (2005) Medial preoptic area interactions with the nucleus accumbens-ventral pallidum circuit and maternal behavior in rats. Behav Brain Res 158:53–68.

O’Connell LA, Hofmann HA (2011) The vertebrate mesolimbic reward system and social behavior network: a comparative synthesis. J Comp Neurol 519:3599–3639.

Olazábal DE, Young LJ (2006) Oxytocin receptors in the nucleus accumbens facilitate “spontaneous” maternal behavior in adult female prairie voles. Neuroscience 141:559–568.

Oz M, Kolaj M, Renaud LP (2001) Electrophysiological evidence for vasopressin V(1) receptors on neonatal motoneurons, premotor and other ventral horn neurons. J Neurophysiol 86:1202–1210.

Panksepp J (1981) The ontogeny of play in rats. Dev Psychobiol 14:327–332.

Paul MJ, Terranova JI, Probst CK, Murray EK, Ismail NI, de Vries GJ (2014) Sexually dimorphic role for vasopressin in the development of social play. Front Behav Neurosci 8:58.

Pellis SM, Iwaniuk AN (2000) Comparative analyses of the role of postnatal development on the expression of play fighting. Dev Psychobiol 36:136–147.

Pellis SM, Pellis VC (1987) Play-fighting differs from serious fighting in both target of attack and tactics of fighting in the laboratory ratRattus norvegicus. Aggress Behav 13:227–242.

Pitkow LJ, Sharer CA, Ren X, Insel TR, Terwilliger EF, Young LJ (2001) Facilitation of affiliation and pair-bond formation by vasopressin receptor gene transfer into the ventral forebrain of a monogamous vole. J Neurosci 21:7392–7396.

Pittman QJ, Kombian SB, Mouginot D, Chen X, van Eerdenberg FJ (1998) Electrophysiological studies of neurohypophysial neurons and peptides. Prog Brain Res 119:311–320.

Reppucci CJ, Gergely CK, Veenema AH (2018) Activation patterns of vasopressinergic and oxytocinergic brain regions following social play exposure in juvenile male and female rats. J Neuroendocrinol.

Rood BD, De Vries GJ (2011) Vasopressin innervation of the mouse (Mus musculus) brain and spinal cord. J Comp Neurol 519:2434–2474.

Schneider M, Koch M (2005) Deficient social and play behavior in juvenile and adult rats after neonatal cortical lesion: effects of chronic pubertal cannabinoid treatment. Neuropsychopharmacology 30:944–957.

Seamans JK, Floresco SB, Phillips AG (1998) D1 receptor modulation of hippocampal-prefrontal cortical circuits integrating spatial memory with executive functions in the rat. J Neurosci 18:1613–1621.

Smith CJW, Poehlmann ML, Li S, Ratnaseelan AM, Bredewold R, Veenema AH (2017) Age and sex differences in oxytocin and vasopressin V1a receptor binding densities in the rat brain: focus on the social decision-making network. Brain Struct Funct 222:981–1006.

Smith KS, Berridge KC (2005) The ventral pallidum and hedonic reward: neurochemical maps of sucrose “liking” and food intake. J Neurosci 25:8637–8649.

Stavast CJ, Erkeland SJ (2019) The Non-Canonical Aspects of MicroRNAs: Many Roads to Gene Regulation. Cells 8.

Taylor GT (1980) Fighting in juvenile rats and the ontogeny of agonistic behavior. J Comp Physiol Psychol 94:953–961.

Taylor PV, Veenema AH, Paul MJ, Bredewold R, Isaacs S, de Vries GJ (2012) Sexually dimorphic effects of a prenatal immune challenge on social play and vasopressin expression in juvenile rats. Biol Sex Differ 3:15.

van den Berg CL, Hol T, Van Ree JM, Spruijt BM, Everts H, Koolhaas JM (1999) Play is indispensable for an adequate development of coping with social challenges in the rat. Dev Psychobiol 34:129–138.

van Kerkhof LWM, Damsteegt R, Trezza V, Voorn P, Vanderschuren LJMJ (2013) Social play behavior in adolescent rats is mediated by functional activity in medial prefrontal cortex and striatum. Neuropsychopharmacology 38:1899–1909.

van Kerkhof LWM, Trezza V, Mulder T, Gao P, Voorn P, Vanderschuren LJMJ (2014) Cellular activation in limbic brain systems during social play behaviour in rats. Brain Struct Funct 219:1181–1211.

van Leeuwen FW, Caffe AR, De Vries GJ (1985) Vasopressin cells in the bed nucleus of the stria terminalis of the rat: sex differences and the influence of androgens. Brain Res 325:391–394.

Veenema AH, Bredewold R, De Vries GJ (2012) Vasopressin regulates social recognition in juvenile and adult rats of both sexes, but in sex- and age-specific ways. Horm Behav 61:50–56.

Veenema AH, Bredewold R, De Vries GJ (2013) Sex-specific modulation of juvenile social play by vasopressin. Psychoneuroendocrinology 38:2554–2561.

Veenema AH, Neumann ID (2009) Maternal separation enhances offensive play-fighting, basal corticosterone and hypothalamic vasopressin mRNA expression in juvenile male rats. Psychoneuroendocrinology 34:463–467.

Wang Z, Bullock NA, De Vries GJ (1993) Sexual differentiation of vasopressin projections of the bed nucleus of the stria terminals and medial amygdaloid nucleus in rats. Endocrinology 132:2299–2306.

Williams DJ, Crossman AR, Slater P (1977) The efferent projections of the nucleus accumbens in the rat. Brain Res 130:217–227.

Zahm DS, Heimer L (1990) Two transpallidal pathways originating in the rat nucleus accumbens. J Comp Neurol 302:437–446.

Zhao C, Chang L, Auger AP, Gammie SC, Riters LV (2020) Mu opioid receptors in the medial preoptic area govern social play behavior in adolescent male rats. Genes Brain Behav 19:e12662.

